# Floating EMG Sensors and Stimulators Wirelessly Powered and Operated by Volume Conduction for Networked Neuroprosthetics

**DOI:** 10.1101/2022.03.09.483628

**Authors:** Laura Becerra-Fajardo, Marc Oliver Krob, Jesus Minguillon, Camila Rodrigues, Christine Welsch, Marc Tudela-Pi, Albert Comerma, Filipe Oliveira Barroso, Andreas Schneider, Antoni Ivorra

## Abstract

**Background:** Implantable neuroprostheses consisting of a central electronic unit wired to electrodes benefit thousands of patients worldwide. However, they present limitations that restrict their use. Those limitations, which are more adverse in motor neuroprostheses, mostly arise from their bulkiness and the need to perform complex surgical implantation procedures. Alternatively, it has been proposed the development of distributed networks of intramuscular wireless microsensors and microstimulators that communicate with external systems for analyzing neuromuscular activity and performing stimulation or controlling external devices. This paradigm requires the development of miniaturized implants that can be wirelessly powered and operated by an external system. To accomplish this, we propose a wireless power transfer (WPT) and communications approach based on volume conduction of innocuous high frequency (HF) current bursts. The currents are applied through external textile electrodes and are collected by the wireless devices through two electrodes for powering and bidirectional digital communications. As these devices do not require bulky components for obtaining power, they may have a flexible threadlike conformation, facilitating deep implantation by injection.

**Methods:** We report the design and evaluation of advanced prototypes based on the above approach. The system consists of an external unit, floating semi-implantable devices for sensing and stimulation, and a bidirectional communications protocol. The devices are intended for their future use in acute human trials to demonstrate the distributed paradigm. The technology is assayed *in vitro* using an agar phantom, and *in vivo* in hindlimbs of anesthetized rabbits.

**Results:** The semi-implantable devices were able to power and bidirectionally communicate with the external unit. Using 13 commands modulated in innocuous 3 MHz HF current bursts, the external unit configured the sensing and stimulation parameters, and controlled their execution. Raw EMG was successfully acquired by the wireless devices at 1 ksps.

**Conclusions:** The demonstrated approach overcomes key limitations of existing neuroprostheses, paving the way to the development of distributed flexible threadlike sensors and stimulators. To the best of our knowledge, these devices are the first based on WPT by volume conduction that can work as EMG sensors and as electrical stimulators in a network of wireless devices.

## I. Background

Clinically available neuroprostheses have been demonstrated to improve the quality of life of patients with neurological disorders or injuries, as these individuals gain significant functional improvement [1]. Additionally, some neuroprosthetic technologies have granted the possibility to perform intuitive control of robotic prostheses as they create an interface between the machine and the patient’s nervous system [2].

In the framework of the EXTEND collaborative project, funded by the European Commission and in which the authors participate, it has been defined the concept of Bidirectional Hyper-Connected Neural Systems (BHNS). The BHNS concept refers to systems consisting of minimally invasive communication links between multiple nerves or muscles in the body and external devices which may be interconnected between them (Fig. 1). The concept requires the deployment of dense networks of wireless active implantable medical devices (AIMDs) that can perform distributed electrical stimulation and sensing. These wireless AIMDs must communicate in real time with external devices and tools that process and analyze the neuromuscular activity and control the stimulation and the action of machines. BHNS could, for example, be used for 1) tremor management in essential tremor and Parkinson’s disease by tremor prediction using electromyography (EMG) [3], and subsequent intramuscular stimulation [4]; 2) performing functional neuromuscular stimulation [5]; or 3) interfacing with assistive wearable robots for spinal cord injury (SCI) and interfacing with prostheses providing control and artificial sensory feedback [6].

**Fig. 1.**
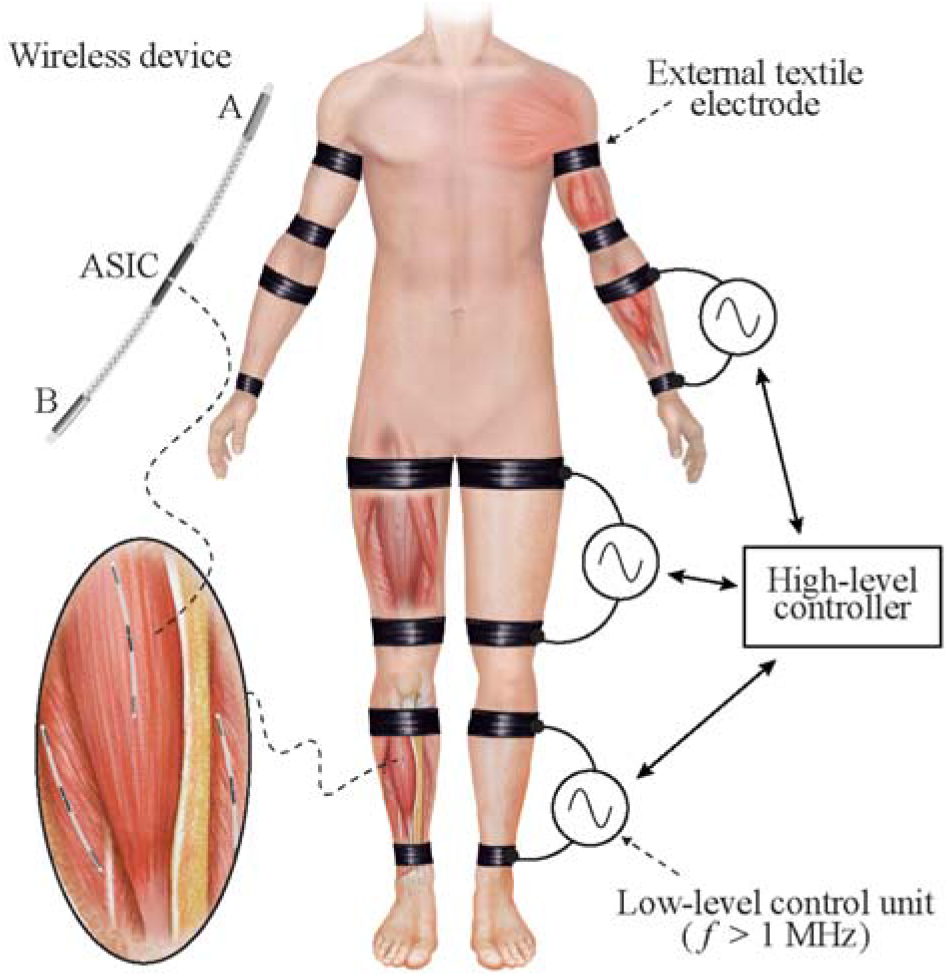
Schematic representation of a Bidirectional Hyper-Connected Neural System – BHNS. High frequency (> 1 MHz) volume conduction is used to wirelessly power and communicate with implantable devices for sensing and stimulation. The high-level controller of the external system communicates with several low-level control units that deliver current bursts through external textile electrodes. The envisioned wireless implantable devices will have a thread-like conformation to facilitate their deployment by injection. These devices will consist of a flexible body with two electrodes at their opposite ends (Electrodes ‘A’ and ‘B’) and their electronics will be integrated in an ASIC.

Most neuroprostheses for chronic use, which are required to accomplish the BHNS concept, are implantable systems that consist of a central unit electrically connected to electrodes at target sites (e.g., cuff electrodes on peripheral nerves) by means of leads (i.e., wires) [7,8]. However, these neuroprostheses present limitations that restrict their use. Those limitations, which are most detrimental in the case of motor neuroprostheses, often arise from their bulkiness and the need to perform complex surgical implantation procedures because of the leads [1,9]. The leads also tend to fail due to mechanical stress, particularly in the case of motor neuroprostheses because electrodes are commonly deployed in long and mobile anatomical regions. As an alternative to these AIMDs based on central units, for the case of motor neuroprostheses, it was proposed and assayed the development of distributed networks of wireless intramuscular microstimulators that integrate the electronics and the electrodes [10,11]. The devices communicate with external systems to perform neuromuscular stimulation aiming at motor restoration in patients suffering from motor paralysis [12]. The microstimulators were percutaneously implanted via injection, thereby avoiding complex surgeries. However, the devices were stiff and considerably large (diameters > 2 mm), making them unsuitable for their use in a dense network of wireless microstimulators.

The form factor of these wireless devices is mostly limited by the method used to power them. Batteries, with their intrinsic limited lifespan and large volume [13], are not suitable as primary sources of power in very small electronic implants. These floating implants, in clinical use or preclinically demonstrated, are typically powered by means of wireless power transfer (WPT) methods such as inductive coupling - the most established WPT method - [14–17], ultrasonic acoustic coupling [18–20] and capacitive coupling [21–23]. These methods have obtained high miniaturization levels, at the expense of link efficiency, penetration depth, or functionality. Recent reviews on these methods can be found in [13,24–26].

In the last years we have proposed and demonstrated very thin microstimulators whose operation is based on rectification of volume conducted innocuous high frequency (HF) current bursts to cause local low frequency currents capable of stimulation [27,28]. A portion of the HF current picked up by the devices is not directly rectified to perform stimulation but used to power the electronics of the implant (e.g., for control and communications) and, in this sense, it can be understood that the implants employ WPT based on volume conduction. In fact, in a series of recent works, we have advocated for, and studied, the use of volume conducted HF current bursts applied through textile electrodes to power elongated implants in general, not only stimulators [29–31]. Remarkably, according to the approach we propose, the implants can be conceived as thin, flexible and elongated devices suitable for implantation by means of injection [27]. The approach uses the body as an electrical conductor of innocuous and imperceptible HF current bursts that are applied by an external system using external textile electrodes. The implants do not require bulky components within their body to be electrically fed, allowing the integration of the electronics in an application-specific integrated circuit (ASIC). They pick up a small portion of the energy available through their two electrodes located at the opposite ends of the implant. The energy is used to power and bidirectionally communicate with the external system (Fig. 1). The elongated and ultrathin form factor allows the deployment of multiple implants in a single body region, favoring their use in dense networks of wireless AIMDs, such as those required by BHNS.

In the general architecture of BHNS, the external system will consist of one top-level controller that communicates with the wireless AIMDs through several external low-level control units that act as bidirectional gateways (i.e., protocol translators) between the implants and the external controller (Fig. 1). The low-level units apply bursts of HF currents to power and communicate with the wireless devices.

In Becerra-Fajardo et al. [27] we reported the development and *in vivo* evaluation of semi-rigid, thin and addressable stimulators based on this WPT approach. The injectable devices were made only of off-the-shelf components, and had an overall diameter of 2 mm. Although successfully demonstrated in animals, due to their invasiveness, these devices are not adequate to conduct assays in humans. Therefore, to be able to acutely demonstrate in humans the feasibility of the BHNS concept, within the framework of the EXTEND project it was decided to develop semi-implantable devices that consist of ultrathin and short intramuscular electrodes, made with thin-film technology, that are connected to a miniature electronic circuit to be fixed on the skin. Not only the electronics of the devices were upgraded for EMG sensing [32] but they were also upgraded for communication capabilities. Here we report the development and evaluation of these semi-implantable devices and the external system that powers and controls them. To the best of our knowledge, the wireless devices presented here are the only devices based on WPT by volume conduction that can work both as EMG sensors and as electrical stimulators to form networks of wireless devices.

## II. Methods

### 1. Developed system

The two fundamental parts of the developed system are 1) the external system that delivers the HF current bursts for wireless powering and bidirectional communications from the external system to the floating semi-implantable devices (i.e., downlink), and from these devices to the external system (i.e., uplink), and 2) the floating devices. These floating devices are semi-implantable devices composed of intramuscular electrodes connected to a miniature external electronic circuit (Fig. 2). The external system, through the two external textile electrodes of a specific low-level control unit, delivers HF current bursts that power the floating devices located between the textile electrodes. Using the same HF current bursts, the control unit sends commands to the wireless devices to operate them, and, depending on the command, delivers HF currents that are modulated by the wireless devices to generate an uplink reply.

**Fig. 2.**
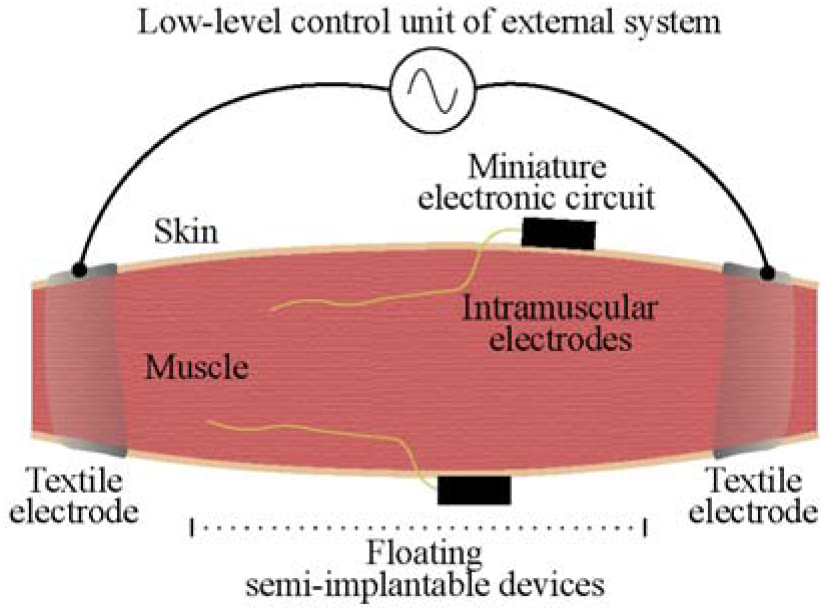
Schematic representation of the floating semi-implantable devices used for acute implantation, and a low-level control unit of the external system. The semi-implantable devices include injectable intramuscular electrodes and a miniature electronic circuit adhered to the skin. The low-level unit of the external system delivers HF current bursts for powering, EMG sensing and electrical stimulation.

#### 1.1. External system

##### 1.1.1. Hardware

The external system integrates a low-power PC/104 single board computer (CMA34CRQ2100HR by RTD Embedded Technologies, Inc) acting as a high-level controller that can communicate with several low-level units. The basic architecture of the low-level control units is shown in Fig. 3. The units include a small single-board computer - hereinafter “digital unit” - (Raspberry Pi 4 2G Model B, by Raspberry Pi Foundation) that controls the delivery of the HF current bursts and the bidirectional communications. They also include a HF generator and modulator (4064 by B&K Precision, Corp.) connected to a custom-made power amplifier consisting of five high voltage amplification modules, based on a high speed, high voltage operational amplifier (ADA4870 by Analog Devices, Inc.), which are connected in series through transformers. The output of the power amplifier is connected to a sensing resistor, and to a pair of textile electrodes attached to the skin. The sensing resistor, acting as a shunt resistor, is used by a custom-made demodulator for measuring the current flowing through the tissues. This current is monitored by the digital unit, especially during uplink, when the floating devices modulate their current consumption, creating minute variations in the current. Fig. 3 shows a schematic representation of the modulating signal and resulting modulated signal for downlink and power bursts (blue waveforms), the voltage obtained across the sensing resistor, which corresponds to a downlink and an uplink section, and the corresponding demodulated uplink data (red waveforms). For illustration purposes, the uplink modulation index used in the scheme is 10%, higher than the < 1% modulation index of the load modulation performed by the floating devices. The load modulation scheme is explained below.

**Fig. 3.**
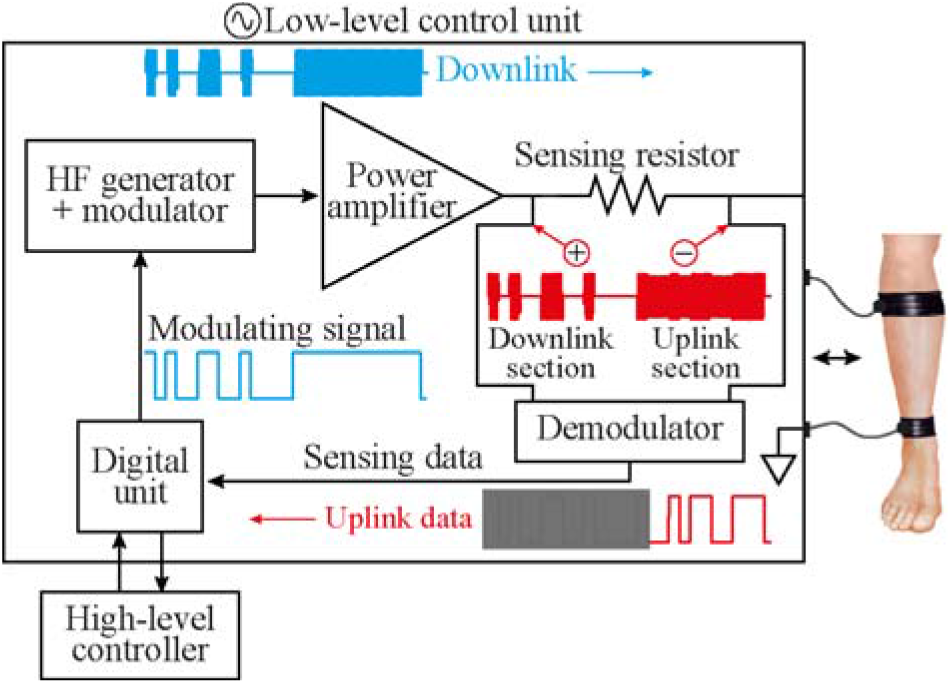
Basic architecture of the low-level control unit. The unit is governed by the high-level controller and communicates with the wireless devices located in the tissue between the two external electrodes of the control unit. The device includes a digital unit that generates a modulating signal for a HF generator and modulator. The modulated HF current - carrying downlink information and bursts of power - is amplified and applied to the tissues through the external textile electrodes (blue waveforms). This same current is continuously monitored by the digital unit through a sensing resistor and a demodulator, to detect and decode minute current variations (< 1 %) generated when the wireless devices do uplink (red waveforms).

For experimental convenience, in the present study the reported assays were performed without using the PC/104 single board computer as the top control device: the Raspberry Pi 4 acting as the digital unit was commanded through the Ethernet interface from a computer running Matlab (R2019b, by Mathworks, Inc). This computer acted as the high-level controller.

Through its universal asynchronous receiver-transmitter (UART) port, the digital unit of the low-level unit (i.e., the Raspberry Pi 4) generates an amplitude-shift keying (ASK) signal to modulate the HF current bursts for downlink communications and for powering the wireless devices. For uplink communications, the digital unit reads the information amplified and filtered by the demodulator using the UART interface. For both downlink and uplink, the information is sent at a rate of 256 kbps. This implies that each byte has a duration of 39.06 μs (1 start bit + 1 byte + 1 stop bit).

##### 1.1.2. HF bursts for power

The external system delivers an initial long HF sinusoidal burst coined “Power up”. This 30 ms burst is required to power up the wireless devices located between the two external electrodes. After this, the wireless devices are kept energized, and running in an idle mode, by delivering short bursts with a repetition frequency (*F*) of 50 Hz, and a duration (*B*) of 1.6 ms. As later explained, the EMG analog front-end (AFE) of the wireless devices saturates during the burst. The repetition frequency and duration of these bursts is selected to 1) minimize their impact in the EMG front-end, favoring faster recovery from saturation and longer windows in-between bursts that can be used for EMG acquisition; 2) obtain a duty cycle (*D*) that is low enough to avoid tissue heating, and 3) deliver enough energy to keep the semi-implantable devices powered.

##### 1.1.3. Digital communications

A communication protocol stack structured in layers and based on the Open System Interconnection (OSI) model, has been created for performing the bidirectional communications between the external system and the wireless devices. The protocol was created to ensure that the minimum required data frames were used in the downlink and uplink, minimizing the time the HF bursts are active. The active time of the HF bursts is related with the specific absorption rate (SAR), which is limited by safety standards (explained below).

The application layer implemented in this protocol stack includes 13 different downlink commands to control (e.g., configure EMG acquisition and stimulation) and interrogate (e.g., ping, get samples) the wireless devices. The frames that encapsulate the commands and the replies used in the bidirectional communications are encoded employing Manchester coding, which offers two advantages: has no dc component to avoid charge injection when applying the HF current bursts, and it provides a first error detection mechanism. Other additional error detection mechanisms are parity bit, frame length and command code.

A detailed description of the process performed by the external unit to command the wireless devices, request replies, and the timings used to deliver the HF current bursts is included in Supplementary Methods, Supplementary file 1. All downlink commands and uplink replies are preceded by one initialization byte. That is, each frame consists of one initialization byte followed by one or more bytes containing the information. All uplink replies are preceded by a downlink command. Between one downlink command (e.g., Get sample) and the corresponding uplink reply (e.g., Send sample) there is a period of 2.3 ms in which no bursts are delivered by the external system. This time is required for processing purposes inside the wireless device.

Supplementary Table 1, Supplementary file 1, reports the downlink commands included in the communication protocol stack, as well as the estimated transmission time required for them. The protocol allows to control up to 256 wireless devices located between two external electrodes. To avoid replicating the instructions to several wireless devices, the protocol enables the use of 256 groups of devices that can be configured simultaneously, or that can be requested to do a specific function (e.g., start sensing or stop sensing). This minimizes the delivery of HF bursts for downlink and avoids misusing the powering/communication channel.

The stimulation and sensing configuration payloads of the communication protocol stack are reported in Supplementary Table 2, Supplementary file 1.

Supplementary Table 3, Supplementary file 1, reports the uplink replies, description and timings used by the wireless devices to send information to the external system. The replies either correspond to 1) an acknowledge (ACK), 2) a sample or 3) the configuration information currently defined in the wireless device. When the external system sends a “Get configuration” request, the uplink frame is constructed using the same format as in the downlink. In the case of “Get sample” and “Retry sample”, the device replies with a frame consisting of 10 bits corresponding to the sample, and 2 more bits corresponding to an internal counter, which is used to control the correct uplink of samples.

##### 1.1.4. Compliance with electrical safety standards

ICNIRP and IEEE standards define safety levels with respect to human exposure to electromagnetic fields. These standards protect against health effects related to tissue heating [33,34]. Heating limits are expressed by the standards in terms of the SAR, which can be related to the electric field at a point as:

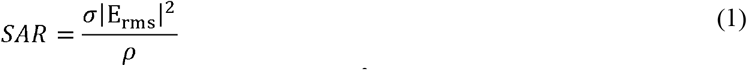

where *σ* is the conductivity of the tissue (S/m), *ρ* is the tissue density (kg/m^3^), and E is the electric field strength in tissue (V/m) averaged over 6 minutes for local exposure. By applying the HF sinusoidal currents in the form of short bursts, the applied E_rms_ is:

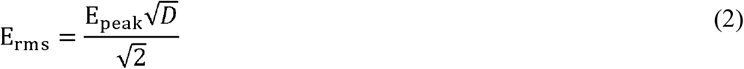

where E_peak_ is the applied peak electric field (V/m), and *D* is the duty cycle. In the case of periodic bursts (frequency *F*; duration *B*), as is the case of the power maintenance bursts, *D* is equal to *F·B*.

Equation (2) allows to calculate the applied E_rms_ and SAR in terms of the Power up time, and the types of functions requested by the external system, including the commands sent in the downlink, and the information replied in the uplink. Supplementary Tables 1 and 3, Supplementary file 1, can be used to calculate the duty cycle corresponding to the commands used in a bidirectional sequence. The duty cycle can be calculated as:

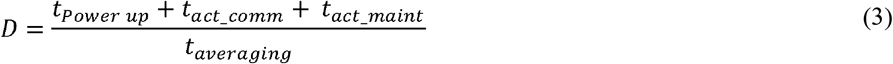

where *t_Power up_* is the Power up time (i.e., 30 ms), *t_act_comm_* corresponds to the active time of the commands, both downlink and uplink, *t_act_maint_* corresponds to the active time of the power maintenance bursts (which have a duty cycle equal to *F·B*), and *t_averaging_* is the averaging time of the rms (360 s, equivalent to the 6 minutes averaging time established for the SAR calculation). The duration of the power maintenance bursts is equivalent to *t_averaging_* minus the total time used for Power up and bidirectional commands.

In the case of human limbs, the standards define a maximum SAR of 20 W/kg for persons in restricted environments at frequencies below 6 GHz, averaged over 10 g cube of tissue [33,34].

#### 1.2. Wireless devices for EMG sensing and electrical stimulation

The wireless devices are composed of thin and flexible intramuscular electrodes made in thin-film technology, a miniature electronic circuit with off-the-shelf components, and an intermediate printed circuit board (PCB) that connects both parts.

##### 1.2.1. Intramuscular electrodes

###### 1.2.1.1. Active sites design

The intramuscular electrodes are based on similar thin-film electrodes reported in [35], which are intended for acute implantation. To define the geometry of the electrode contacts (i.e., the active sites, the actual electrodes strictly speaking), simulations were performed to test the ability of the electrodes to 1) obtain enough power to supply the wireless circuit for continuous EMG recording, and 2) generate stimulation pulses with amplitudes above 2 mA and below 4 mA. Other functions performed by the circuit were not evaluated as they require less power than that needed during EMG acquisition and electrical stimulation. The simulations were performed by modeling the tissues surrounding the intramuscular electrodes and the presence of the electric field applied by the external system using a Thévenin equivalent circuit [29]. See Supplementary Methods, Supplementary file 1, for details of the numerical methods for the intramuscular electrodes design.

Besides power supply and current delivery capabilities, another aspect that was considered in the geometrical design of the active sites was the charge injection limit. If the applied pulses exceed the maximum charge injection capacity of the electrodes, irreversible reactions may occur, which can cause damage both to the electrodes and to the tissues. In the case of smooth platinum, the charge injection capacity is defined in the range between 50 and 150 μC/cm^2^ for 200 μs, biphasic, charge-balanced pulses [36]. In the case of microrough platinum, the charge injection capacity increases to 500 μC/cm^2^ [37].

###### 1.2.1.2. Final conformation

The final geometry of the electrodes is based on the results obtained with finite element method (FEM) simulations and with circuit simulations performed with SPICE. Both are described in Supplementary Methods, Supplementary file 1. The polyimide filament that contains the electrode contacts and the tracks has a width of 0.42 mm, a length of 81.6 mm, and a thickness of 0.02 mm (Fig. 4 (a)). Between the centers of the contacts there is a distance of 30 mm, and the contacts have a length of 7.5 mm and a width of 0.265 mm (Fig. 4 (b)). The edges of the contacts are rounded to avoid sharp corners that would lead to high current densities. The four electrode contacts are located on both faces of the filament: 2 contacts in the top layer, and 2 contacts in the bottom layer. The distal contacts (top and bottom) are electrically connected in the miniature electronic circuit, to use them as a single distal electrode with a total surface area of 3.8 mm^2^. This electrode, coined Electrode ‘A’ (Fig. 4 (a)), is the stimulation electrode when the semi-implantable device is in stimulation mode. The two proximal contacts (top and bottom) are also short-circuited, to use them as a single proximal electrode (Electrode ‘B’, return electrode in stimulation mode) with a total surface area of 3.8 mm^2^. Electrodes ‘A’ and ‘B’ are used as the inputs for the EMG analog front-end (AFE) and are indirectly used by the load modulator during uplink. Both circuits are explained below.

**Fig. 4.**
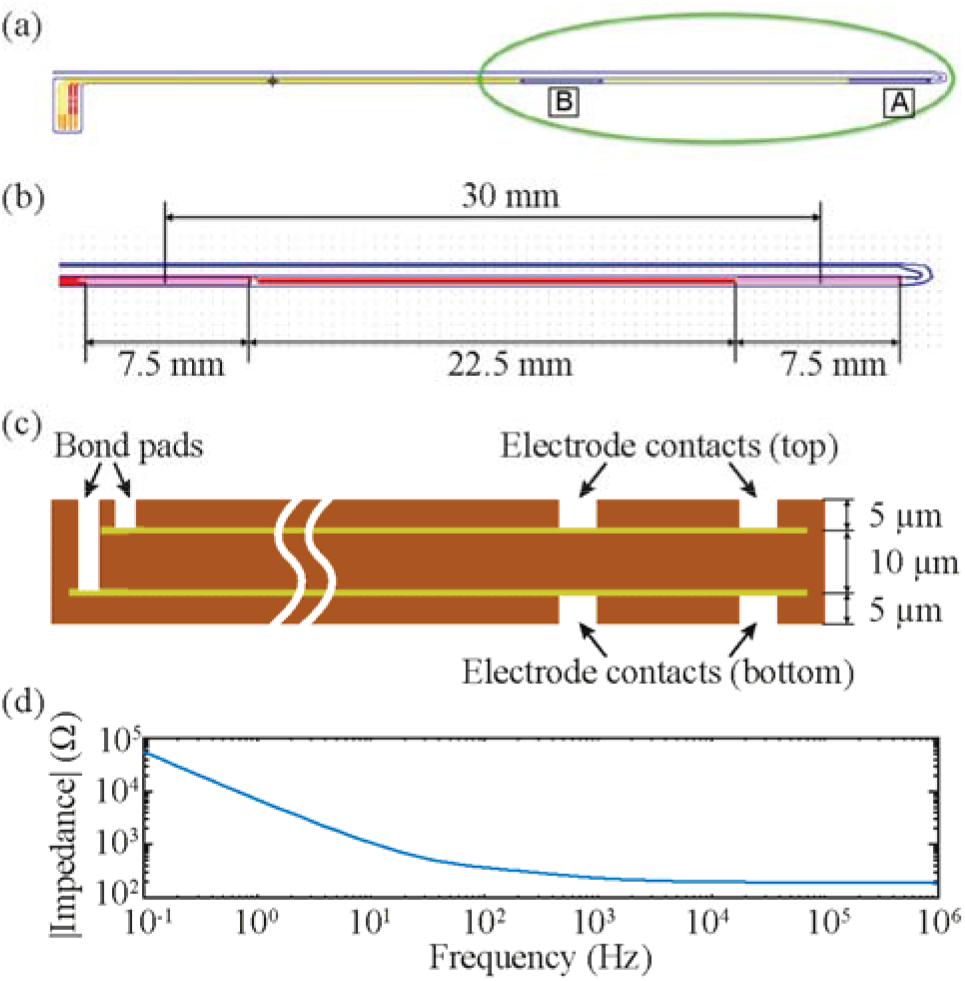
Intramuscular electrodes. (a) Top view of the electrodes showing a distal contact (Electrode ‘A’), a proximal contact (Electrode ‘B’), and the bonding pads of the four electrode contacts. (b) Detail of the area where the distal and proximal electrodes are located. (c) Cross section of the intramuscular electrodes (not to scale). Gold and platinum are used as metallization layers for the electrode tracks and contacts (600 nm thick per layer), and polyimide is used as substrate and cover material. (d) Impedance measurement of the intramuscular electrodes in a bipolar arrangement (short circuited distal contacts against short circuited proximal contacts) in a 0.9% NaCl solution.

Fig. 4 (c) shows the construction of the intramuscular electrodes. In essence it consists of a polyimide substrate (thickness: 10 μm) with a patterned metallization on both sides and a thin polyimide coating (thickness: 5 μm) on both sides with openings for the active sites and the bonding pads. The metallization is uniform, that is, it is the same metallization both for the active sites and for the connection tracks from these to the bonding pads. A relatively thick layer of gold (450 nm) beneath the platinum layer (150 nm) wires ensures high conductivity for the tracks. Microrough platinum coating is used to increase the surface area of the electrodes, therefore increasing their charge injection capacity and improving power transfer.

As the proximal end of the thin-film electrodes is connected to the external miniature electronic circuit, they have the same implantation limitation as those used in [35,38]: the intramuscular electrodes cannot be implanted directly using a hypodermic needle as introducer, because the circuit would prevent the complete extraction of the needle after insertion. To overcome this drawback, the distal end of the filament containing the four electrode contacts has a U-shape structure with a guiding filament (Fig. 4 (b)) that facilitates the insertion of the electrodes in the muscle. The main filament with the active sites runs externally to the needle, while the thin guiding filament is inserted through the lumen of a 23 G hypodermic needle (Sterican 4665600 by B. Braun Melsungen AG) having a length of 60 mm and an outer diameter of 0.6 mm. The end of the guiding filament is adhered to the Luer lock of the needle. To reduce the risk that the needle bevel cuts the guiding filament during implantation, the bevel is smoothed with a laser (Picco Laser, by O.R. Lasertechnologie, Germany) prior to inserting the guiding filament inside the needle.

The small intermediate PCB is used for 1) doing the electroplating process to coat the electrode contacts with microrough platinum [39], and 2) connecting the injectable electrodes to the miniature electronic circuit. The polyimide electrodes are bonded to the PCB using the Microflex technology to achieve an electrical and mechanical connection [40]. After electroplating, the circuit of the semi-implantable device is stacked on top of the small PCB via surface-mount vertical headers, and the final electronic assembly is protected within a 3D printed housing (Fig. **5**).

**Fig. 5.**
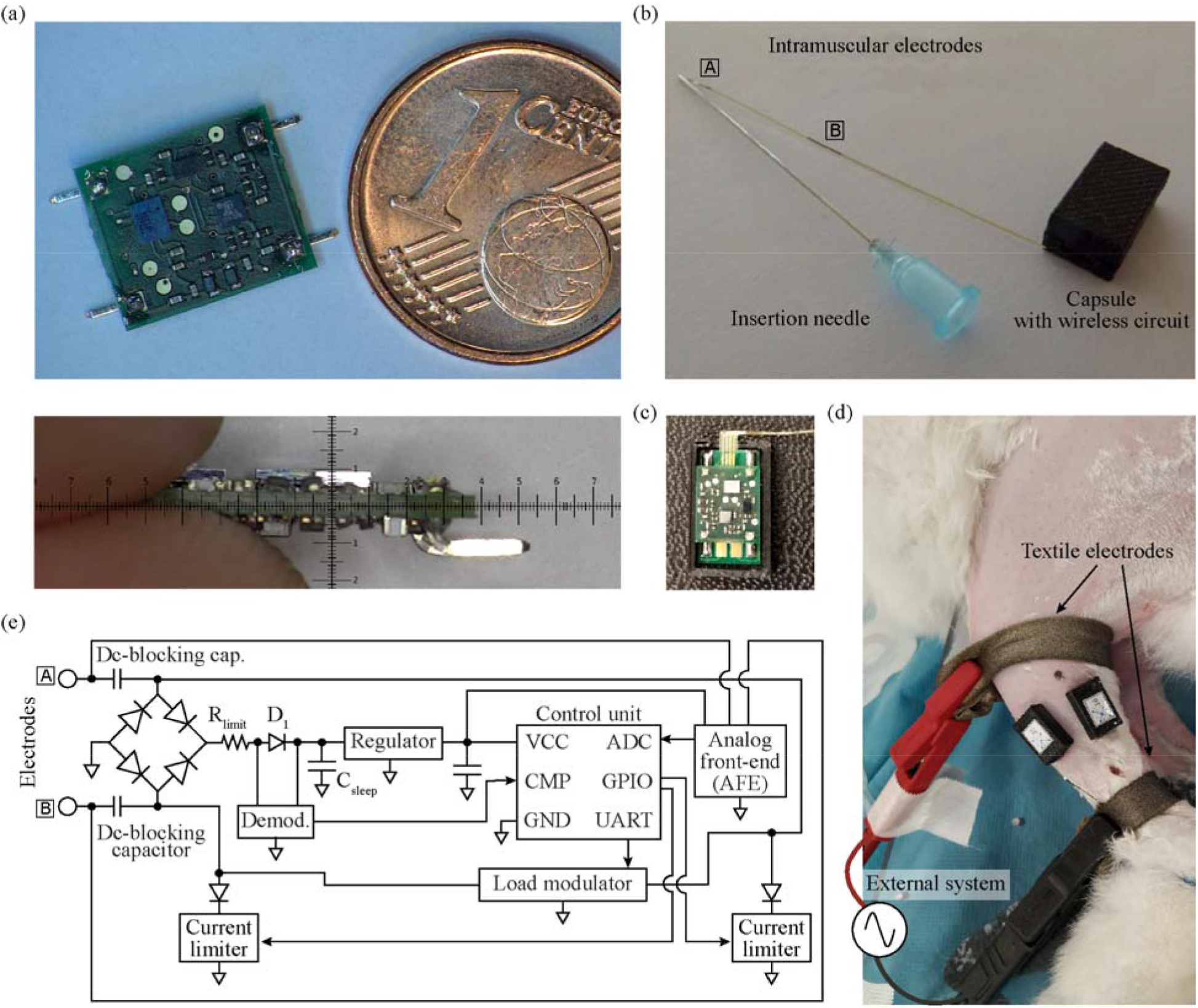
Wireless device composed of intramuscular electrodes and miniature electronic circuit. (a) Top: circuit board (length: 10 mm; width: 8.5 mm), with four headers for their connection to the 4 electrode contacts of the intramuscular electrodes. Bottom: lateral view of the circuit (height: 2.4 mm). (b) Final conformation of the wireless device, including intramuscular electrodes and capsule protecting circuit. The guiding filament of the electrodes is inserted in the lumen of the insertion needle. (c) Circuit housed in a polymer capsule, connected to the intramuscular electrodes using a small PCB. (d) Image of rabbit’s hindlimb with two wireless devices implanted in the tibialis anterior and gastrocnemius medialis muscles. (e) Basic architecture of the wireless circuit shown in (a).

##### 1.2.2. Miniature electronic circuit

The electronic architecture of the wireless device used for sensing and stimulation and which is powered and operated by HF volume conduction is partially based on [27]. Yet the architecture presented here has several advantages over the design reported in the past, including better power efficiency, bidirectional communications with higher data rates, the integration of an AFE for EMG acquisition, an ultra-low power microcontroller with several peripherals and running a finite-state machine, and a communication protocol stack (described above) for a more robust control from the external system.

The electronic circuit has a length of 10 mm, and a width of 8 mm. The surface-mount vertical headers shown in Fig. 5 (a), connect the circuit to the intermediate PCB that includes the intramuscular electrodes. The final conformation of the wireless device, including the insertion needle and the capsule to protect the wireless circuit, is shown in Fig. 5 (b).

###### 1.2.2.1. Power regulation and dc-blocking

Fig. 5 (e) shows the basic architecture of the electronic circuit. The distal and proximal electrodes of the polyimide filament are connected to the electronic circuit through two dc-blocking capacitors, which are aimed to prevent dc currents that can damage both the tissues and the electrodes [41]. A bridge rectifier based on four Schottky diodes (RB521ZS-30 by ROHM Co., Ltd.) provides full-wave rectification for the HF current picked up by the intramuscular electrodes. A limiting resistor followed by a Schottky diode (D_1_), and a set of three 2.2 μF capacitors connected in parallel (C_sleep_) provide a smoothed input for a low-dropout linear regulator (ADP7112ACBZ-2.5 by Analog Devices, Inc.). The output of this regulator is connected to a set of two 10 μF capacitors connected in parallel. This provides a stable dc voltage for the control unit and the rest of the electronics during and in-between HF bursts. Sets of capacitors connected in parallel are used instead of single capacitors to ensure a proper capacitance and voltage rating in a miniature surface-mount package.

###### 1.2.2.2. Control unit

The wireless electronic circuit is controlled by an ultra-low power microcontroller (MKL03Z32CAF4R by NXP Semiconductors N.V.) with an Arm Cortex M0+ core. This 2 mm × 1.6 mm microcontroller is larger than that of [27], but includes several peripherals required by the circuit, a much lower power-up time, several low-power modes, and more general-purpose input/output (GPIO) pins. The integrated high-speed comparator is used as the final stage of the downlink demodulator (explained below), and its output is connected to the integrated low-power UART receiver for further decoding. The UART transmitter generates the modulating signal for the load modulator used for uplink communications. Two digital outputs of the GPIO pins are used to control the switches of the current limiters for electrical stimulation, and the analog-to-digital converter (ADC) is used as the last stage of the AFE. Both circuits are described below. Finally, a low-power timer is used to control the sampling times during sensing, and to control the time-outs of the software.

The control unit is configured to have three types of power consumption states: 1) idle mode, when the implant is waiting for commands from the external system, 2) processing and basic operations mode, when the implant decodes and processes the information coming from the external system and performs uplink communications and stimulation, and 3) sensing mode, when the implant is acquiring EMG activity and most of the peripherals of the microcontroller are powered. The change of state is defined by the information received from the external system during downlink. The control unit is usually kept in a very-low power standby state (i.e., idle mode) to maintain the wireless circuit energized, avoiding the need to deliver a long Power up HF burst to power-up the microcontroller in-between bursts. Since the circuit lacks an active power source as a battery, it naturally shuts down at any time by disabling the bursts of HF current applied by the external system.

The initialization byte sent by the external system is used by the control unit as the source for a hardware-based automatic wake-up matching, to wake up from the idle mode and go into the processing and basic operations mode. In the case the byte received matches the wake-up code already defined in the firmware, the wireless device wakes up and waits for the next byte with information to decode. The information includes the address of the wireless device or group of devices, and the command sent to it.

###### 1.2.2.3. Downlink demodulation and uplink modulation

For downlink, a demodulator was designed consisting of two voltage dividers whose outputs are connected to the inputs of a high-speed comparator integrated in the control unit. The output of the first voltage divider corresponds to the envelope of the downlink signal modulated in the HF current burst. The second voltage divider, which is connected immediately after the Schottky diode D_1_, behaves with the set of 2.2 μF capacitors as a low-pass filter, creating a threshold proportional to the output of the first voltage divider. In other words, even if the amplitude obtained across the intramuscular electrodes varies due to fluctuations in the electric field applied by the external system, the outputs of both voltage dividers maintain their proportionality. The downlink signal is used for 1) receiving data at a rate of 256 kbps or 2) receiving triggers to wake up the control unit using the synchronization byte or perform a function as uplink communications.

The uplink is based on load modulation using on-off keying. The external system applies a HF current burst, which is detected by the wireless devices to do their specific uplink sequence. Supplementary Methods, Supplementary file 1 describes the uplink sequence as well as the timings required for each type of reply (e.g., ACK, send configuration, send sample). The floating device identifies this burst, starting the uplink section shown in Fig. 3. Its control unit generates a 256 kbps Manchester-encoded modulating signal with its UART transmitter. This modulating signal is used to command a set of transistor switches that short-circuit the inputs of the full-wave rectifier (Fig. 5 (e)). This results in virtually short-circuiting Electrodes ‘A’ and ‘B’, thus modulating the current consumption of the wireless device. These minute variations in consumption are seen in the sensing resistor of the external system during the uplink section (Fig. 3, red waveforms), and are then demodulated and decoded by the digital unit. Load modulation is completely innocuous to the tissue: there is no electrical stimulation generated when uplink is performed. As a risk mitigation strategy, the load modulator is connected to the intramuscular electrodes through the dc-blocking capacitors.

###### 1.2.2.4. EMG analog front-end (AFE)

One of the most challenging characteristics of the wireless device proposed here is the use of only two electrodes (Electrodes ‘A’ and ‘B’) for powering, bidirectional communications, electrical stimulation, and EMG sensing. As the wireless device powers from the volume-conducted alternating HF currents, very little consumption imbalances during the positive and negative semicycles may slightly charge the electrodes during these episodes, saturating the EMG amplifier. For this reason, the input of the EMG amplifier avoids the use of high dc-blocking capacitances that could prolong such saturation beyond the burst. Fig. 6 shows the basic architecture of the designed EMG amplifier. A four-resistor difference amplifier with a gain of 18.4 dB was designed based on a micropower operational amplifier (ADA4505 by Analog Devices, Inc). The amplifier is biased using a split resistor (R_F_/2), and a capacitor is added to each amplifier input, to act with the input resistors (R_IN_) as a first low-pass filter. A Zener diode clipper is included in both inputs of the operational amplifier for protection. The difference amplifier is followed by cascaded active band-pass filters with a bandwidth of 1 kHz. The overall gain of the AFE is 54 dB, and its output is connected to the ADC of the control unit, which is configured at a resolution of 10 bits.

**Fig. 6.**
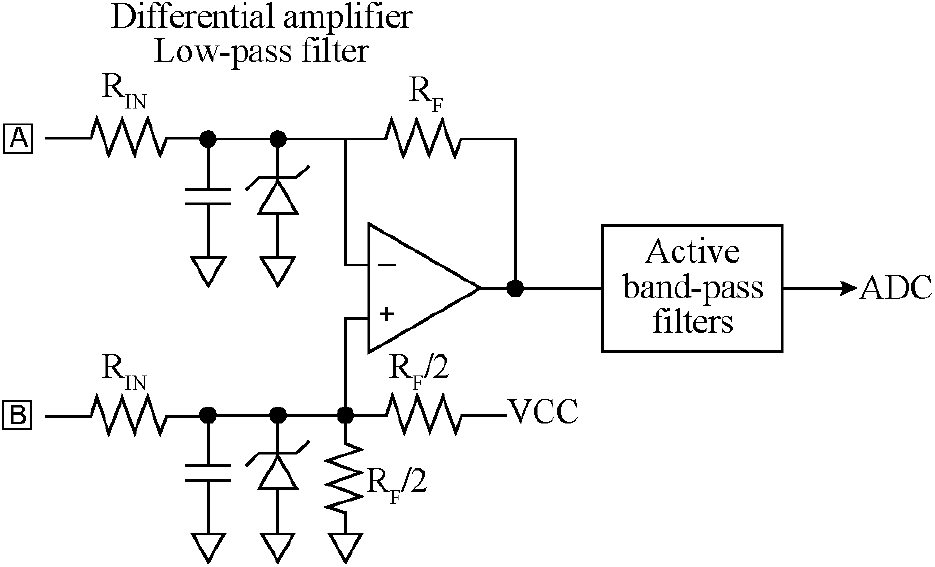
Basic architecture of the AFE designed for the wireless device. The input of the AFE is the same pair of electrodes used for powering, communication and electrical stimulation

During EMG sensing, the control unit is constantly monitoring when a HF burst is applied by the external system using the high-speed comparator. When this happens, depending on the sampling frequency set by the external system, the control unit replaces the samples corresponding to this saturation with a constant value (e.g., 5 samples for a sampling frequency of 1 ksps). The number of samples to replace is implicitly set by the external system when configuring the sampling rate of the sensing mode. The replacement of the samples facilitates to identify the moments in which the AFE was saturated when the complete recording is uploaded to the external system, without misinterpreting the saturation with high amplitude EMG activity.

###### 1.2.2.5. Electrical stimulation

As mentioned above, the electrical stimulation mechanism of the wireless devices is based on the rectification of the volume conducted HF current bursts to cause local low frequency currents capable of stimulation. This is performed using two independent current limiters, each one connected to a Schottky diode (RB521ZS-30 by ROHM Co., Ltd.) that is connected to an electrode (Electrode ‘A’ or ‘B’) by means of a dc-blocking capacitor (Fig. 5 (e)). The architecture of the current limiter is explained in depth in the Supplementary Methods, Supplementary file 1. Each current limiter is connected/disconnected from the load (i.e., the tissue) using a switch based on a N-channel MOSFET (DMN2990UFZ by Diodes Incorporated) controlled by the control unit of the wireless device through one GPIO.

Using the communication protocol explained above, the external system configures the control unit to deliver either monophasic or biphasic stimulation pulses and the polarity of the pulse or pulses. The pulse width, number of stimulation pulses and frequency of stimulation are set by the external system using specific timings of the HF current bursts. The wireless device detects these bursts using its demodulator, and the control unit activates the current limiters accordingly. In the case of biphasic pulses, an interphase dwell is generated by the external system, which is automatically detected by the wireless device. In both monophasic and biphasic configurations, 60 μs after the stimulation sequence (i.e., a monophasic or biphasic pulse) is completed, a short HF current burst of 800 μs is applied by the external system. This short burst is used by the wireless device to short-circuit the dc-blocking capacitors, clearing the charge that has accumulated in them in case there is a mismatch between the current limiters. In this way, passive charge balance is accomplished. Supplementary Methods, Supplementary file 1 shows a schematic representation of the activation of the current limiters according to the stimulation configuration defined by the external unit, and the resulting low-frequency contents of the current flowing through the wireless device.

### 2. *In vitro* validation

To evaluate the impedance of the intramuscular electrodes, a bipolar arrangement was implemented, in which the short circuited distal contacts (Electrode ‘A’) were measured against the short circuited proximal contacts (Electrode ‘B’). An impedance measurement system (Autolab PGSTAT302N by Deutsche METROHM GmbH & Co. KG) was employed using this arrangement in a 0.9% NaCl solution. In addition, in a dry setup, the resistance between the electrode contacts and the PCB pads was measured using a hand-held multimeter (38XR-A by Amprobe Instrument Corporation).

The *in vitro* model used for the evaluation of the wireless device consisted in a 6.5 cm diameter agar cylinder made from a NaCl solution with a conductivity of 0.57 S/m, equivalent to that of muscle tissue at 3 MHz (Fig. **7** (a)) [42]. The agar cylinder was cut longitudinally, and one side was placed on a 3D printed structure to ensure stability. The intramuscular electrodes were placed in a position such that the wireless circuit could lie just by the cylinder (Fig. 7 (b)). After carefully placing the second half of the cylinder over the electrodes, the agar phantom was held together with the use of two aluminum external electrodes (width: 1.5 cm), which were strapped around the cylinder at a distance of 12 cm. The aluminum electrodes were connected to the external system using high strength alligator clips (AK 2 S, by Hirschmann Test and Measurement).

**Fig. 7.**
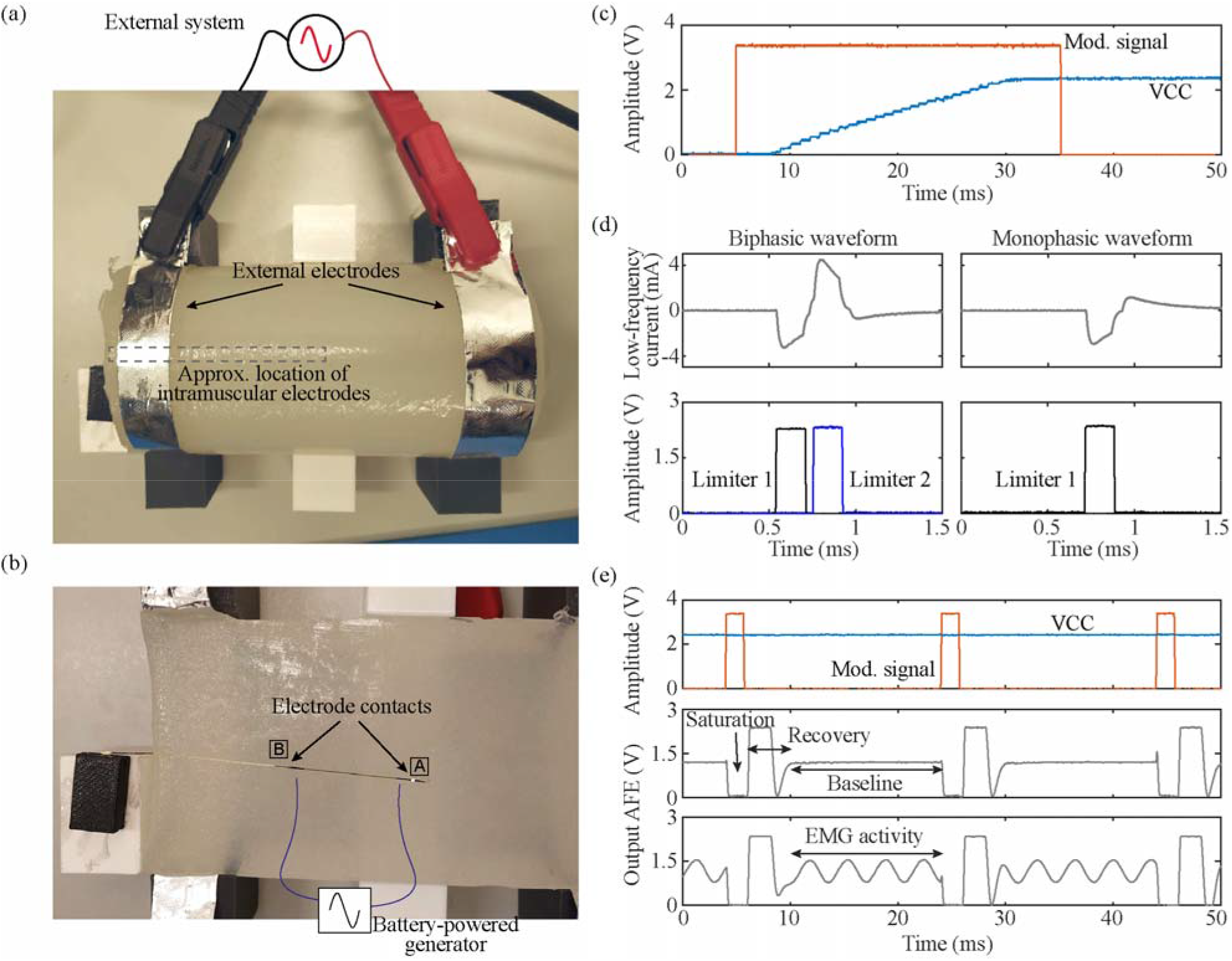
*In vitro* setup and results. (a) Complete *in vitro* setup. (b) Detail of half the agar cylinder, showing the position of the intramuscular electrodes and the wireless electronic circuit. The approximate location of the electrode wires connected to the battery-powered generator for EMG testing is shown in blue. (c) Modulating signal of the external system (Mod. signal), and output of the wireless device’s regulator (VCC), which is used to power the circuit’s electronics. A ‘high’ of the modulating signal corresponds to the delivery of HF currents. (d) Filtered biphasic and monophasic stimulation waveforms obtained when the wireless device rectifies the HF current immediately after the external system triggers a stimulation. The current limiters of the circuit are enabled/disabled by the control unit to obtain such waveforms. (e) Output of the AFE of the wireless device measured with an oscilloscope when no sinusoidal signal is present, and when there is. The AFE saturates due to the HF currents applied by the external system (see corresponding modulating signal), and recovers to have a stable state (baseline) for EMG acquisition. VCC is stable even if the HF current is applied in the form of bursts, and the circuit is at its maximum power consumption.

For performing electrical measurements, the miniature wireless electronic circuit was replaced by a larger evaluation board that included test points. In the case of stimulation, the proximal electrode (Electrode ‘B’) of the circuit was connected to the parallel combination of a 10 Ω resistor and a 2.2 μF capacitor. This measurement circuit acts as a low-pass filter (LPF) (cutoff frequency: 7.2 kHz) for the current that flows through the intramuscular electrodes, facilitating recording the low-frequency contents of the current flowing through the wireless device. All the measurements were performed using a battery-powered oscilloscope (TPS2014 by Tektronix, Inc.), except for the current consumption of the circuit, which was measured using a battery-powered multimeter (38XR-A by Amprobe Instrument Corporation). The signals obtained with the oscilloscope were smoothed in Matlab (R2019b, by Mathworks, Inc) using a centered moving average filter with a window length of 5 samples.

To test the capability of the wireless circuit to acquire EMG, a low frequency potential dipole was created in the saline agar cylinder by connecting a battery-powered sinusoidal generator (72-505 by Tenma Test Equipment) to two kynar-insulated silver plated copper wire electrodes (exposed length: 3.65 mm, diameter: 0.25 mm). The wire electrodes were inserted into the cylinder with a separation distance of approximately 30 mm between them, and were placed very close to the electrode contacts of the intramuscular electrodes. A schematic representation of this generator and the approximate location of the wire electrodes is shown in Fig. 7 (b). The generator was set to apply sinusoidal waveforms with amplitudes and frequencies similar to those expected for EMG (maximum amplitude: 1 mV; frequency range: 20 – 420 Hz in 16 steps defined by the generator).

### 3. *In vivo* validation

#### 3.1. Animal handling

The animal procedure was approved by the Ethical Committee for Animal Research of the Centre for Comparative Medicine and Bioimage (CMCiB) of the Germans Trias i Pujol Research Institute (IGTP), and by the Catalan Government (project number: 10109). Three New Zealand White male rabbits with a mass from 4 kg to 4.4 kg were employed for the proof-of-concept, one per session. The first session was meant for training and preliminary testing the devices; no results from it are reported here. The third session was meant for evaluating the implantation and explantation procedures of the intramuscular electrodes; no results are reported here.

For sedation and initial anesthesia, dexmedetomidine (0.15 mL/kg), butorfanol (0.04 mL/kg) and ketamine (0.08 mL/kg) were intramuscularly administered between 15 to 30 minutes prior to the preparation of the animal. Then, the left hindlimb of the rabbit was shaved, from the head of the femur to the mid tarsus. In addition, a depilatory cream (Veet sensitive skin, by Reckitt Benckiser Group plc) was applied for 3 minutes, the limb was thoroughly rinsed, and cleaned with alcohol and Betadine using a sterile gauze. The animal was transferred to an anesthetic circuit using endotracheal intubation and anesthesia was maintained with isoflurane (2-3% and 0.5% of O_2_). Ringer’s lactate (8 mL/h) was administered intravenously, and the animal was constantly monitored with a capnograph, pulse oximeter, esophageal temperature sensor and non-invasive blood pressure sensor. A heating pad and fluid warmer were employed during the entire session. At the end of the study, the animal was euthanized with an overdose of pentobarbital sodium (10 mL).

#### 3.2. Implantation procedure and extraction

One semi-implantable device was used in the tibialis anterior (TA) muscle (Fig. 5 (b)) and a second semi-implantable device was used in the gastrocnemius medialis (GA) muscle, of the left hindlimb of the rabbits. Target muscles were identified by palpation and the approximate site for deploying the intramuscular electrode (i.e., close to the motor point) was located using anatomical cues. To verify the correct location of this target, a hook wire intramuscular electrode (003-400160-24 by SGM d.o.o.) was inserted into the muscle, and an Ag/AgCl gel electrode (model 2228 by 3M) placed on the thigh of the animal was used as a return electrode. Both electrodes were connected to a commercial current generator (Ultra 20, by Kegel8), which delivered 1 to 2 mA biphasic pulses (pulse width: 200 μs) at 100 Hz. If the induced movement was considered weak or did not match the expected joint movement, a new position was defined, and a hook wire electrode was inserted to verify the response. After the approximate location of the target point was successfully obtained, it was identified with a marker, and the hook wire was extracted.

The needle with the thin-film intramuscular electrodes was longitudinally introduced from the hock to the identified location close to the motor point, resulting in an insertion length of about 5 cm. The housing of the circuit was adhered to the skin of the rabbit to avoid the accidental extraction of the electrodes. Finally, the end of the guiding filament adhered to the Luer lock was cut, and the needle was gently extracted.

After the *in vivo* assays were finished, the intramuscular electrodes were extracted by holding the guiding filament and the electrodes’ filament and gently pulling out.

#### 3.3. Stimulation assays

After implantation, the animal was laid sideways and the left hindlimb was placed on a force test bench to measure isometric plantarflexion and dorsiflexion forces using a load cell (STC-10kgAL-S by Vishay Precision Group, Inc.). The output of the load cell was connected to a custom-made signal conditioning system composed of a precision instrumentation amplifier with a voltage gain of 100 V/V (LT1101 by Analog Devices, Inc.), a RC low-pass filter (cutoff frequency: 2 kHz), and an isolation amplifier with a fixed gain of 8.2 V/V (ACPL-790B by Broadcom, Corp.). This filter was used to avoid noise and HF interference that could be coupled to the test bench through the animal limb and the load cell. Its cutoff frequency was low enough to attenuate the HF current (3 MHz), and the harmonics created by the downlink modulating signal. The resulting signal of the isolation amplifier was recorded using a 10-bit ADC, at a sampling rate of 4 kHz, and was digitally filtered using a 4th-order zero-phase forward and reverse Butterworth low-pass filter, with a cut-off frequency of 10 Hz. The load cell was fixed to a custom-made acrylic board, the hock was fixed to the board using an atraumatic padded clamp, and the foot was tied with cable ties to the load cell (Fig. 8 (a)). The external electrodes consisted in 1.5 cm wide bands made from silver-based stretchable conductive fabric (Shieldex® Technik-tex P130 + B (1150902130TB) by Statex Produktions und Vertriebs GmbH) strapped around the hindlimb of the animal (Fig. 5 (d)). The shortest distance between centers of external electrodes, corresponding to the posterior side of the hindlimb, was 6.5 cm. The electrodes were connected to the external system using high strength alligator clips (AK 2 S, by Hirschmann Test and Measurement). The HF current bursts applied were monitored using a differential active probe (TA043 by Pico Technology) and a current probe (TCP2020 by Tektronix, Inc.) connected to a battery-powered oscilloscope (TPS2014 by Tektronix, Inc.).

**Fig. 8.**
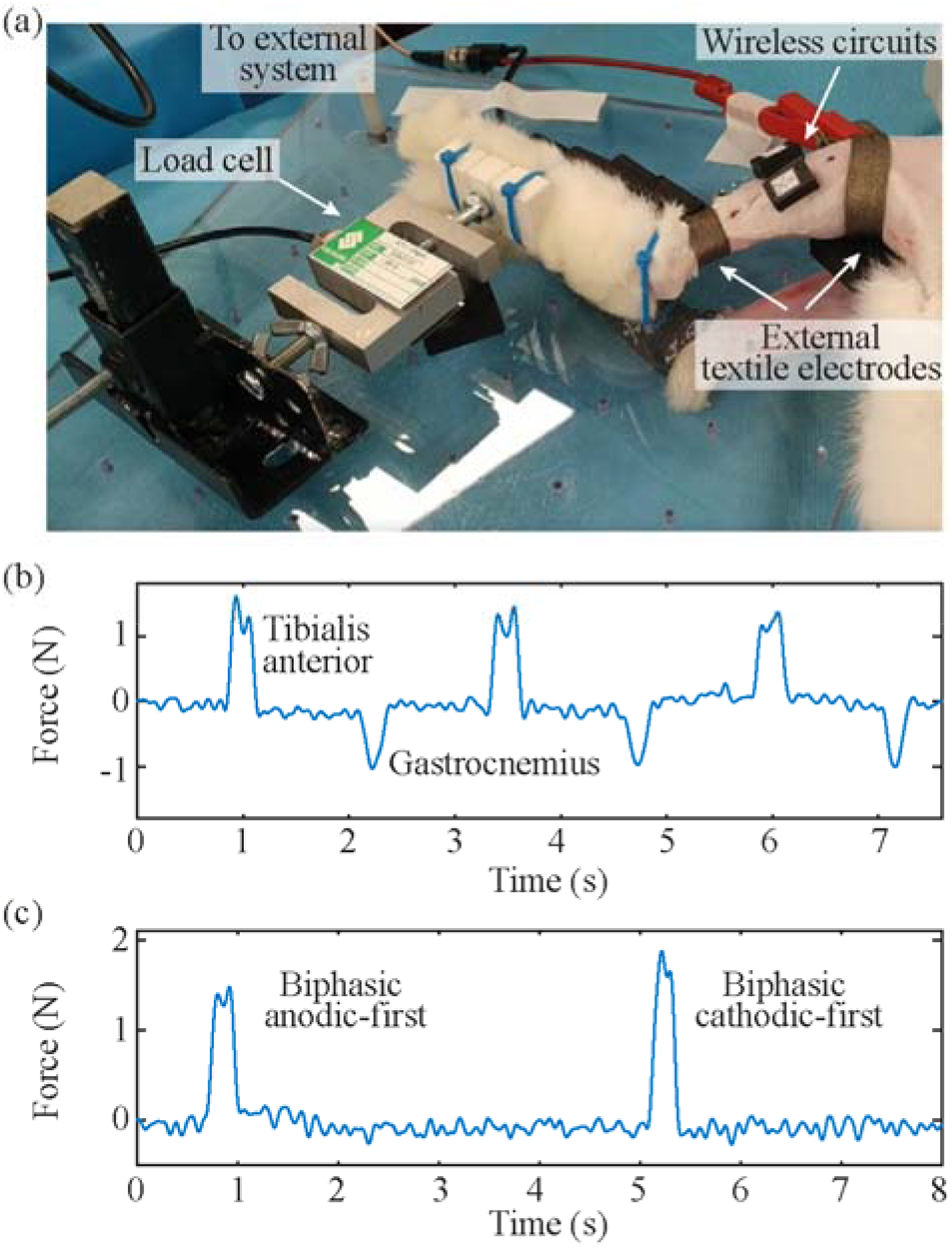
*In vivo* assays for electrical stimulation. (a) Setup used for recording forces during electrical stimulation. Positive forces correspond to plantaflexion movements, while negative forces correspond to dorsiflexion movements. (b) Forces obtained by addressing, using the external system, the devices deployed in TA and GA muscles, both configured as biphasic, anodic-first stimulation. (c) Forces obtained after the external system changed the internal stimulation configuration (anodic/cathodic pulse first stimulation) of a device deployed in TA muscle. Both stimulation sequences were done with 20 stimulation pulses at 100 Hz with a pulse width of 100 μs.

#### 3.4. EMG sensing assays

Muscle activity was triggered with the pedal withdrawal reflex: a deep nociceptive reflex [43], which contracts the flexor muscles and relaxes the extensors muscles. To do so, the level of anesthesia was decreased until a reflex was obtained by pinching with forceps the interdigital space of the hindlimb [44]. When the pinch was done, an EMG acquisition was triggered by sending from the external system the start and stop sensing functions to the group of semi-implantable devices. Once the reflex was performed, the deep plane of anesthesia was recovered by increasing the isoflurane rate or, alternatively, in two instances, by intravenous injection of a low dose of propofol (0.25 mg/kg). During the assays, it was verified that the TA muscle contracted with the withdrawal reflex, as it is the case in humans [45], but as the level of anesthesia was low, immediately after the reflex the GA muscle could be also activated. To avoid loosening the external electrodes, therefore affecting the coupling for volume conduction, the foot of the rabbit during EMG sensing assays was not allowed to move by holding it with the force test bench used during electrical stimulation assays (Fig. 8 (a)).

## III. Results

### 1. *In vitro* assays

For the *in vitro* assays, the external system was configured to deliver HF bursts with an amplitude of 39 V, a sinusoidal current frequency of 3 MHz, a burst duration (*B*) of 1.6 ms, and a burst frequency (*F*) of 50 Hz. This amplitude was the minimum required for the correct operation of the wireless device. The combination of duration and repetition frequency of the power maintenance bursts was enough to keep the wireless devices powered after the Power up stage (i.e., idle mode).

#### 1.1 Powering the wireless device

The impedance magnitude of the intramuscular electrodes, as measured across two electrode pairs in 0.9 % NaCl (Fig. 4 (d)), is flat beyond 10 kHz, and lower than 190 Ω at 1 MHz. This indicates that, at high frequencies, the impedance of the electrode-electrolyte interface is negligible in comparison with the combined resistance of the saline solution and of the tracks. Thus, at high frequencies, the contact impedance between the tissue and the circuit terminals will mostly correspond to the resistance of the tracks. The resistance measured between the distal electrode contacts (Electrode ‘A’) and the PCB pad was 56 Ω; while that measured between the proximal electrode contacts (Electrode ‘B’) and the PCB pad was 28 Ω.

As expected by design, the wireless electronic circuit was successfully powered through the intramuscular electrodes using the HF current bursts delivered by the external system.

The time required by the control unit to initialize was much shorter than that required in [27], decreasing the duration of the initial Power up burst from 85 ms to 30 ms, therefore lowering the applied SAR. Fig. 7 (c) shows the modulating signal of the external system (i.e., when 3.3 V are measured at “Mod. signal”, the HF current is delivered), and how the regulator of the wireless circuit is able to provide a stable output (VCC) of 2.3 V approximately 26 ms after the modulating signal is activated. The stability of this VCC is required for the correct behavior of the circuit, specially the AFE.

The current consumptions during the idle mode, during the processing and basic operations mode and during the sensing mode were measured in the *in vitro* setup (Fig. 7 (a)) and the evaluation board connected to the intramuscular electrodes. The circuit required 180 μA, 185 μA, and 400 μA respectively to power all its electronic components and the control unit’s peripherals required for the power mode defined.

#### 1.2 Electrical stimulation

The external system was able to set the electrical stimulation configuration (biphasic/monophasic stimulation, anodic/cathodic pulse first) using the communication protocol designed for the bidirectional communications. This configuration can be changed at any time by sending a new configuration from the external system. The pulse duration, number of pulses and frequency of the pulses were successfully managed by the external system by defining the duration, number and frequency of the bursts of HF current.

Fig. 7 (d) shows an example of a biphasic, cathodic-first waveform, and a monophasic, cathodic-first waveform obtained in the *in vitro* setup. As indicated above, this signal is obtained after low-pass filtering the partially rectified current produced by the circuit, and digitally filtering it. The current limiters are activated using the digital outputs of the control unit, when the device decodes that a specific HF burst is meant for electrical stimulation (see the activation signal for the limiters in the lower graphs of Fig. 7 (d)). During these stimulation bursts, the amplitude of the HF current was increased to 47 V to deliver more energy to the semi-implantable devices, obtaining a maximum stimulation amplitude of 4.5 mA.

In the case of the biphasic waveforms, even if the circuit has mismatches between the current limiters that define the cathodic and anodic pulses, the dc-blocking capacitors and its discharge mechanism can balance the charge applied in each pulse. In the case of monophasic waveforms, this same protection mechanism is used to perform purely passive charge-balanced stimulation [41].

#### 1.3 EMG acquisition

The floating sinusoidal generator was set to apply different frequencies and amplitudes, which were then acquired by the wireless electronic circuit connected to the intramuscular electrodes. Even though the highest consumption of the control unit was during EMG sensing, the circuit was able to power and keep a steady voltage of 2.3 V during the sensing process (Fig. 7 (e), top, VCC). The output of the AFE measured from a test point of the evaluation board is shown in Fig. 7 (e). During the application of the HF bursts, the AFE saturated, but the circuit successfully recovered and returned to its baseline state to acquire EMG activity in-between bursts (Fig. 7 (e), middle). The time window available to do a proper EMG acquisition was approximately 14 ms (see baseline time in Fig. 7 (e), middle). When the sinusoidal generator was turned on, the AFE was able to filter and amplify the signal picked up by the intramuscular electrodes. This signal was then digitized using the ADC of the control unit. Fig. 7 (e), bottom, shows an example of the output of the AFE when the sinusoidal generator was delivering a waveform of 280 Hz with an amplitude of 800 μV.

### 2. *In vivo* assays

The intramuscular electrodes were easily inserted into the TA and GA muscles of the rabbits using the dedicated 23 G hypodermic needle. The insertion mechanism was robust and minimally invasive. Supplementary file 2 shows a video with the complete implantation procedure. Fig. 5 (d) shows an image of two wireless devices implanted in the hindlimb of the animal, and the location of the textile electrodes that connect to the low-level control unit of the external system. At the end of the assays, the electrodes were safely removed.

For these assays, the external system was configured to deliver HF bursts with an amplitude of 52 V, a frequency of 3 MHz, a burst duration (*B*) of 1.6 ms, and a burst frequency (*F*) of 50 Hz. The amplitude of the current during the bursts was 0.3 A. If the external system applies power maintenance bursts, the duty cycle (3) is equal to 0.08 (i.e., *F·B*). This low duty cycle scales down the applied HF bursts to 10.4 V_rms_ and 0.06 A_rms_, resulting in an average power of 0.62 W.

#### 2.1 Electrical stimulation

The external system was able to configure the stimulation parameters of the wireless devices and control the devices to perform electrical stimulation. Fig. 8 (b) shows the forces obtained when the external system addressed the device deployed in the TA muscle, or the device deployed in the GA muscle. Both devices were configured to deliver biphasic, anodic-first pulses. The external system defined the stimulation protocol with pulse widths of 100 μs, and 20 pulses. The TA device was commanded to stimulate at 100 Hz, while the GA device was commanded at 150 Hz. These parameters obtained a maximum force of 1.6 N and 1 N respectively.

Fig. 8 (c) shows the effect of changing the internal configuration of the stimulation. The device deployed in the TA muscle was triggered to perform stimulation with fixed parameters: 20 pulses, 100 μs, and 100 Hz. The device was configured at first for biphasic anodic-first stimulation. Few seconds after doing the stimulation, the device was reconfigured to do biphasic cathodic-first stimulation. The maximum force obtained for the first configuration was 1.5 N, while that of the second configuration was 2 N.

#### 2.2 EMG sensing

The wireless devices reported here were capable of registering EMG signals and sending this information wirelessly to the external system. The circuit recorded raw signals according to the configuration set by the external system.

Fig. 9 shows an example of raw EMG signals obtained simultaneously by two wireless devices after the pedal withdrawal reflex was triggered, with the devices deployed in the TA and the GA muscles respectively (1 ksps). The EMG samples were uploaded to the external system using the communication protocol described above. The floating devices were able to acquire samples before, during and after the contraction. The period corresponding to the contraction can be observed as there is an episode in which the amplitude of the EMG increases significantly.

**Fig. 9.**
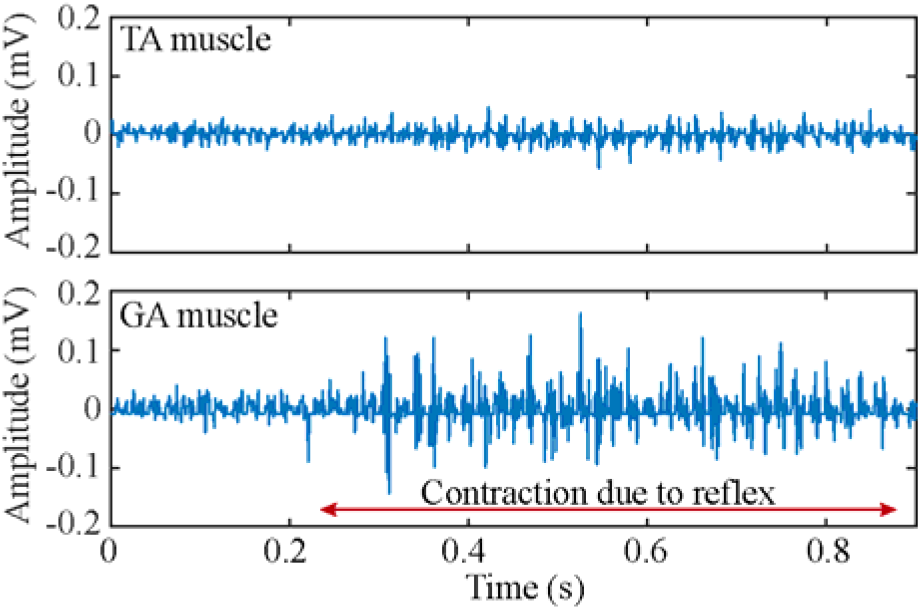
Example of raw EMG obtained after the reflex was triggered using a device implanted in the TA muscle and a second device implanted in the GA muscle. This time window shows the moment after the TA muscle was activated by the reflex and the GA muscle is being activated. The amplitude of the muscular activity during contraction is clearly differentiated from that of the baseline.

#### 2.3 Bidirectional communications

The control unit of the wireless circuit was able to reply to the requests sent from the external system, such as functional tasks (reset, stimulate, start/stop sensing, and send sample), configuration (set/send group belonging, set/send stimulation configuration, and set/send sensing configuration), and uplink special requirements (ping, retry send sample, and acknowledge). The communication protocol supports the possibility of sending these requests to one wireless circuit, or a group of them. This allows the use of the devices as a network of electrical stimulators and sensors, having the possibility to configure all these functionalities from the external system simultaneously.

The commands ‘get sensing configuration’ (downlink) followed by a ‘send sensing configuration’ reply (uplink) have the longer frames of the communication protocol stack. For this reason, they were used to test the quality of the bidirectional communications. Supplementary file 3 shows a screen recording video of a set of 100 ‘get/send sensing configuration’ requests to device 17 deployed in the TA muscle of the rabbit’s hindlimb. The external system sent the request to the wireless devices, and after demodulation, decoding and error detection, device 17 interpreted that it should send the configuration information to the external system. The control unit coded this information and sent it back to the external system (i.e., performed uplink) using the load modulator. The screen shows the information received by the external system after demodulating, decoding, and performing error detection. A success rate of 100% was obtained.

### 3. Compliance with SAR limitation established by electrical safety standards

In the *in vitro* study, the external system was set to apply HF bursts with an amplitude of 39 V, which should produce an electric field in the middle of the region located between the two external textile electrodes with an amplitude of about 325 V/m [30].

The standards indicate that the SAR must be averaged for 6 minutes [33]. In this case, the possible sequences that can be made by the external system to interact with the wireless devices can have several combinations, all starting with a Power up, and having windows of time for power maintenance bursts. Supplementary Table 5, Supplementary file 1, reports three different sequences that could be used in three hypothetical scenarios: 1) Power up and maintenance, 2) Power up, 50 seconds of stimulation (biphasic, 200 Hz, 400 μs pulse width) distributed in 6 minutes, and maintenance; and 3) Power up, 120 cycles composed of 1 s of sensing at 500 sps, 1 s delay for samples uplink and external processing, and 1 s of stimulation (biphasic, 100 Hz, 200 μs pulse width), and maintenance. The second sequence corresponds to the worst-case scenario of stimulation, while the third sequence corresponds to a hypothetical scenario in which neuromuscular activity is used to control stimulation (e.g., for tremor management). To calculate the SAR at a point according to equation (1), the properties of muscle tissue at 3 MHz were used (*σ*: 0.57 S/m [42] and *ρ*: 1090 kg/m^3^ [46]). The SAR obtained for the three sequences was 2.21, 3.40 and 8.10 W/kg respectively, below the 20 W/kg limit established by the standards. The sequences include enough power maintenance time to guarantee a steady power supply in the wireless devices during the second and third sequences (341.06 s and 266.73 s respectively).

## IV. Discussion

In this article, we have reported the design and evaluation of floating devices capable of EMG sensing and electrical stimulation which are wirelessly powered and controlled by an external system using HF volume conduction across living tissues. The assayed devices and systems illustrate the feasibility of forming densely distributed networks of intramuscular wireless microsensors and microstimulators that communicate with external systems for analyzing neuromuscular activity and performing stimulation or controlling external devices.

The main aim of the present study was to test the capabilities of the circuit architecture of the floating devices to obtain power, to bidirectionally communicate with the external system, and to perform EMG sensing and stimulation. To facilitate this evaluation *in vivo*, and to enable future acute human trials, the devices were built as semi-implantable devices: the circuit was designed to be kept outside of the body, and only the ultrathin intramuscular electrodes were to be injected. This semi-injectable conformation is not appropriate for chronic testing due to the possibility of infections, the risk of electrode tear up and accidental extraction. However, the circuit architecture demonstrated here is the basis for the development of an ASIC that includes all its capabilities, and that is intended to be integrated in a flexible threadlike implant with two electrodes at opposite ends. This would allow to have an implantable network of microstimulators and microsensors for BHNS that can be fully configured for specific clinical applications.

One of the main challenges of using the thin-film intramuscular electrodes presented here was the small surface area of the electrodes (3.8 mm^2^) compared to cylindrical electrodes as those assayed in [27] (diameter: 2 mm, length: 3.8 mm; surface area: 27 mm^2^). Despite this limitation, it was possible to accomplish a sufficient surface area to obtain enough electric current to power the wireless miniature electronic circuit, and to perform safe electrical stimulation without exceeding the charge injection capacity of the electrodes.

Another substantial challenge was to combine the low-voltage EMG recordings with the high-voltage HF bursts for power. The WPT approach proposed here requires that the external system continuously delivers bursts of HF current for powering the wireless devices. The wireless device uses the same electrodes for powering, communications, EMG sensing and electrical stimulation. This implies that its AFE saturates when the HF bursts are applied by the external system. Despite this important limitation, and that the presence of the saturation of the AFE is seen in the raw recordings (flat lines in-between the EMG activity), the AFE designed for the circuit was able to have ample time windows (~14 ms) to properly obtain EMG activity, ensuring a fast recovery from the saturation generated with the HF currents. These time windows will allow to record EMG signals with enough quality for their use in the BHNS proposed. Although the EMG recordings obtained were made with the hindlimb held to the load cell setup, it is intended that the system can be used in future acute human trials with free moving individuals, as the textile electrodes can be strapped firmly around the human limbs.

The WPT approach uses a single channel (the tissues) both for powering and communication. This implies that while information is sent from a wireless device to the external system (uplink), other wireless devices cannot send information, nor the external system can do downlink. Another critical aspect of the approach proposed is that the semi-implantable wireless devices use only two electrodes for powering, bidirectional communications, EMG acquisition and electrical stimulation, limiting the possibility of performing several actions simultaneously. However, this characteristic favors the integration of the future ultrathin implantable devices (Fig. 1).

Supplementary Table 6, Supplementary file 1, compares different implantable EMG sensors reported in the literature. Remarkably, all the works identified make use of WPT by inductive coupling. There are two typical conformations: 1) a central unit connected to the electrodes using leads (e.g., the IST-12 [7], the MyoPlant [47], [15], [48] and Ripple [49]), and 2) a cluster of wireless cylindrical implants (IMES [50]). This last conformation is very convenient, as the implantation procedure can be done using minimally invasive techniques [51]. Similarly, we envision our wireless devices as cylindrical and flexible implants that can be deployed by injection. These implants would have a hermetic housing protecting the ASIC with the circuit architecture proposed here. The architecture has successfully demonstrated the possibility to acquire EMG activity, with amplification factor, bandwidth and resolution similar to that of other implantable devices (Supplementary Table 6, Supplementary file 1). This can be useful for applications as neural interfaced assistive wearable robots for SCI, in which the control is done using intramuscular EMG-driven modelling [52]. The architecture has also demonstrated the possibility to perform electrical stimulation, opening the possibility to use the devices in multiple applications, including tremor management in essential tremor and Parkinson’s disease, and prosthetics control with sensory feedback [53].

Another WPT method that is gaining importance due to the possibility of obtaining small form factors is ultrasonic acoustic coupling. Very recently it was demonstrated the possibility of sending information from implantable “motes” to an external system for the use of the implants as neural recorders [54]. Yet the demonstrated implants do not include a hermetic capsule required for long-term implantation [18]. The capsule would increase the size of the mote and could attenuate the ultrasounds, imposing a critical constraint regarding the amount of energy obtained by the implant electronics. Ultrasonic WPT has also demonstrated higher penetration depth compared to inductive and capacitive coupling [24]. However, this is done by means of beam focusing, making it more difficult to arrange the external transmitter for powering and communicating with networks of wireless devices arranged through the body. Another drawback of ultrasonic acoustic coupling is the need to use gel for coupling the external transceiver and the skin [55]. This may cause wounds because of humidity, irritation due to allergens [56], and may be uncomfortable for chronic applications. Our WPT approach based on volume conduction avoids these drawbacks. In [57] it was determined that it would be possible to use existing rechargeable portable batteries (> 100 Wh/kg) for the external system, accomplishing a portable unit that can be easily carried by patients. Also, the textile electrodes proposed do not require the use of gel and can be easily integrated in garments, making it more comfortable for the user, facilitating donning and doffing.

Implant depth is a critical parameter for WPT methods. Here the wireless device was tested *in vitro* by placing the intramuscular electrodes at a depth of 3.25 cm, and it could perform all the functions commanded from the external system. This ideal scenario did not include different conductivity layers as those that could be present due to other tissues. However, we have *in silico* demonstrated using a multilayered geometry, that the electric field generated by the HF current bursts delivered by the external system are coarsely uniform in the region located between the external electrodes [30]. More importantly, we recently demonstrated in arms and lower legs of healthy humans that electric powers above 2 mW and 5 mW respectively could be obtained using needle electrodes (diameter: 0.4 mm, length: 3 mm) implanted approximately 1.75 cm deep [31]. Therefore deeply implanted devices could be powered with this WPT approach.

## V. Conclusions

The present work reports the development and successful evaluation of a technology composed of an external system that powers and controls wireless semi-implantable devices using a WPT approach based on volume conduction. Because the currents applied by the external system were applied in the form of bursts, they were below the heating limits defined by safety standards. The intramuscular electrodes proposed were appropriate for picking up the HF current bursts delivered by the external system to power and operate the designed miniature circuit. The semi-implantable devices for EMG sensing and stimulation could be configured and controlled from the external system using a bidirectional communications protocol that minimizes the application of HF currents. To the best of our knowledge, these are the first wireless devices powered by a WPT approach based on volume conduction that can do electrical stimulation and EMG sensing, and that bidirectionally communicate with the external system. It opens the path to the development of a BHNS that can do distributed electrical stimulation and sensing for neuroprostheses.

## List of abbreviations

BHNS: Bidirectional Hyper-Connected Neural Systems
AIMDs: active implantable medical devices
EMG: electromyography
SCI: spinal cord injury
WPT: wireless power transfer
HF: high frequency
ASICs: application-specific integrated circuits
UART: universal asynchronous receiver-transmitter
ASK: amplitude-shift keying
AFE: analog front-end
OSI: Open System Interconnection
SAR: specific absorption rate
ACK: acknowledge
PCB: printed circuit board
FEM: finite element method
SPICE: Simulation Program with Integrated Circuit Emphasis
GPIO: general-purpose input/output
ADC: analog-to-digital converter
LPF: low-pass filter
CMCiB: Centre for Comparative Medicine and Bioimage
IGTP: Germans Trias i Pujol Research Institute
TA: tibialis anterior
GA: gastrocnemius medialis.

## Declarations

### Authors’ contributions

LBF, JM, AS and AI conceived and designed the study. LBF, JM, CR and AI developed the electronic hardware and the software. MOK, CW, AS and AI designed and developed the intramuscular electrodes. LBF and JM conducted the experiments, performed data analysis and drafted the manuscript. MOK, CR, CW, MTP, AC, FOB, AS and AI revised the manuscript critically. All authors read and approved the final manuscript.

## Acknowledgements

The authors would like to express their gratitude to the team at Centre for Comparative Medicine and Bioimage (CMCiB) of the Gemans Trias i Pujol Research Institute (IGTP) for their work regarding the animal procedures.

## Supplementary information

Supplementary file 1. Appendix: supplementary document with additional information regarding methods and results. This supplementary information is indicated on the body of the research article.

Supplementary file 2. Video with implantation procedure.

Supplementary file 3. Video showing the screenshot of the external system during bidirectional communications in an anesthetized rabbit.

## References

1. Kilgore KL, Anderson KD, Peckham PH. Neuroprosthesis for individuals with spinal cord injury. Neurol Res [Internet]. 2020 Jul 30;1–13. Available from: https://doi.org/10.1080/01616412.2020.1798106

2. Yildiz KA, Shin AY, Kaufman KR. Interfaces with the peripheral nervous system for the control of a neuroprosthetic limb: a review. J Neuroeng Rehabil [Internet]. 2020;17(1):43. Available from: https://doi.org/10.1186/s12984-020-00667-5

3. Basu I, Graupe D, Tuninetti D, Shukla P, Slavin K V, Metman LV, et al. Pathological tremor prediction using surface electromyogram and acceleration: potential use in ‘ON–OFF’demand driven deep brain stimulator design. J Neural Eng. 2013;10(3):36019.

4. Pascual-Valdunciel A, Gonzalez-Sanchez M, Muceli S, Adan-Barrientos B, Escobar-Segura V, Perez-Sanchez JR, et al. Intramuscular stimulation of muscle afferents attains prolonged tremor reduction in essential tremor patients. IEEE Trans Biomed Eng. 2020;1.

5. Hunt AJ, Odle BM, Lombardo LM, Audu ML, Triolo RJ. Reactive stepping with functional neuromuscular stimulation in response to forward-directed perturbations. J Neuroeng Rehabil [Internet]. 2017;14(1):54. Available from: https://doi.org/10.1186/s12984-017-0266-6

6. Sensinger JW, Dosen S. A Review of Sensory Feedback in Upper-Limb Prostheses From the Perspective of Human Motor Control [Internet]. Vol. 14, Frontiers in Neuroscience. 2020. p. 345. Available from: https://www.frontiersin.org/article/10.3389/fnins.2020.00345

7. Hart RL, Bhadra N, Montague FW, Kilgore KL, Peckham PH. Design and Testing of an Advanced Implantable Neuroprosthesis With Myoelectric Control. IEEE Trans Neural Syst Rehabil Eng [Internet]. 2011 Feb [cited 2018 Apr 18];19(1):45–53. Available from: http://ieeexplore.ieee.org/document/5585827/

8. Bergmeister KD, Hader M, Lewis S, Russold M-F, Schiestl M, Manzano-Szalai K, et al. Prosthesis Control with an Implantable Multichannel Wireless Electromyography System for High-Level Amputees. Plast Reconstr Surg. 2016 Jan;137(1):153–62.

9. Memberg WD, Polasek KH, Hart RL, Bryden AM, Kilgore KL, Nemunaitis GA, et al. Implanted Neuroprosthesis for Restoring Arm and Hand Function in People With High Level Tetraplegia. Vol. 95, Archives of Physical Medicine and Rehabilitation. 2014. p. 1201–1211.e1.

10. Kane MJ, Breen PP, Quondamatteo F, ÓLaighin G. BION microstimulators: A case study in the engineering of an electronic implantable medical device. Med Eng Phys. 2011;33(1):7–16.

11. Merrill DR, Lockhart J, Troyk PR, Weir RF, Hankin DL. Development of an Implantable Myoelectric Sensor for Advanced Prosthesis Control. Artif Organs [Internet]. 2011 Mar [cited 2017 Jun 7];35(3):249–52. Available from: http://doi.wiley.com/10.1111/j.1525-1594.2011.01219.x

12. Schulman JH. The Feasible FES System: Battery Powered BION Stimulator. Proc IEEE. 2008;96(7):1226–39.

13. Dinis H, Mendes PM. A comprehensive review of powering methods used in state-of-the-art miniaturized implantable electronic devices. Biosens Bioelectron [Internet]. 2021;172:112781. Available from: https://www.sciencedirect.com/science/article/pii/S0956566320307685

14. Farnsworth BD, Triolo RJ, Young DJ. Wireless implantable EMG sensing microsystem. In: 2008 IEEE Sensors [Internet]. IEEE; 2008 [cited 2017 Sep 8]. p. 1245–8. Available from: http://ieeexplore.ieee.org/document/4716669/

15. Ng KA, Rusly A, Gammad GGL, Le N, Liu S-C, Leong K-W, et al. A 3-Mbps, 802.11g-Based EMG Recording System With Fully Implantable 5-Electrode EMGxbrk Acquisition Device. IEEE Trans Biomed Circuits Syst. 2020;14(4):889–902.

16. Lee J, Leung V, Lee A-H, Huang J, Asbeck P, Mercier PP, et al. Neural recording and stimulation using wireless networks of microimplants. Nat Electron. 2021;4(8):604–14.

17. Loeb GE, Peck RA, Moore WH, Hood K. BION™ system for distributed neural prosthetic interfaces. Med Eng Phys. 2001;23(1):9–18.

18. Piech DK, Johnson BC, Shen K, Ghanbari MM, Li KY, Neely RM, et al. A wireless millimetre-scale implantable neural stimulator with ultrasonically powered bidirectional communication. Nat Biomed Eng [Internet]. 2020;4(2):207–22. Available from: https://doi.org/10.1038/s41551-020-0518-9

19. Chang TC, Weber MJ, Charthad J, Baltsavias S, Arbabian A. End-to-End Design of Efficient Ultrasonic Power Links for Scaling Towards Submillimeter Implantable Receivers. IEEE Trans Biomed Circuits Syst. 2018;12(5):1100–11.

20. Sonmezoglu S, Fineman JR, Maltepe E, Maharbiz MM. Monitoring deep-tissue oxygenation with a millimeter-scale ultrasonic implant. Nat Biotechnol [Internet]. 2021;39(7):855–64. Available from: https://doi.org/10.1038/s41587-021-00866-y

21. Sedehi R, Budgett D, Jiang J, Ziyi X, Dai X, Hu AP, et al. A Wireless Power Method for Deeply Implanted Biomedical Devices via Capacitively Coupled Conductive Power Transfer. IEEE Trans Power Electron. 2021;36(2):1870–82.

22. Sodagar AM, Amiri P. Capacitive coupling for power and data telemetry to implantable biomedical microsystems. In: 2009 4th International IEEE/EMBS Conference on Neural Engineering. 2009. p. 411–4.

23. Hossain ANMS, Erfani R, Mohseni P, Lavasani HM. On the Non-idealities of a Capacitive Link for Wireless Power Transfer to Biomedical Implants. IEEE Trans Biomed Circuits Syst. 2021;15(2):314–25.

24. Barbruni GL, Ros PM, Demarchi D, Carrara S, Ghezzi D. Miniaturised Wireless Power Transfer Systems for Neurostimulation: A Review. IEEE Trans Biomed Circuits Syst. 2020;1.

25. Turner BL, Senevirathne S, Kilgour K, McArt D, Biggs M, Menegatti S, et al. Ultrasound Powered Implants: A Critical Review of Piezoelectric Material Selection and Applications. Adv Healthc Mater. 2021;10(17):2100986.

26. Agarwal K, Jegadeesan R, Guo Y-X, Thakor N V. Wireless Power Transfer Strategies for Implantable Bioelectronics. IEEE Rev Biomed Eng [Internet]. 2017;10:136–61. Available from: http://ieeexplore.ieee.org/document/7879807/

27. Becerra-Fajardo L, Schmidbauer M, Ivorra A. Demonstration of 2-mm-Thick Microcontrolled Injectable Stimulators Based on Rectification of High Frequency Current Bursts. IEEE Trans Neural Syst Rehabil Eng [Internet]. 2017;25(8):1343–52. Available from: http://ieeexplore.ieee.org/document/7726054/

28. Ivorra A, Becerra-Fajardo L, Castellví Q. In vivo demonstration of injectable microstimulators based on charge-balanced rectification of epidermically applied currents. J Neural Eng [Internet]. 2015;12(6):66010. Available from: http://stacks.iop.org/1741-2552/12/i=6/a=066010

29. Tudela-Pi M, Becerra-Fajardo L, García-Moreno A, Minguillon J, Ivorra A. Power Transfer by Volume Conduction: In Vitro Validated Analytical Models Predict DC Powers above 1 mW in Injectable Implants. IEEE Access. 2020;1.

30. Tudela-Pi M, Minguillon J, Becerra-Fajardo L, Ivorra A. Volume Conduction for Powering Deeply Implanted Networks of Wireless Injectable Medical Devices: A Numerical Parametric Analysis. IEEE Access. 2021;9:100594–605.

31. Minguillon J, Tudela-Pi M, Becerra-Fajardo L, Perera-Bel E, Del-Ama AJ, Gil-Agudo Á, et al. Powering electronic implants by high frequency volume conduction: in human validation. bioRxiv [Internet]. 2021 Jan 1;2021.03.15.435404. Available from: https://www.biorxiv.org/content/10.1101/2021.03.15.435404v4

32. Becerra-Fajardo L, Ivorra A. First steps towards an implantable electromyography (EMG) sensor powered and controlled by galvanic coupling. In: World Congress on Medical Physics and Biomedical Engineering 2018 IFMBE Proceedings. Springer Verlag; 2019. p. 19–22.

33. IEEE Standard for Safety Levels with Respect to Human Exposure to Electric, Magnetic, and Electromagnetic Fields, 0 Hz to 300 GHz. IEEE; 2019.

34. International Commission on Non-Ionizing Radiation (ICNIRP). Guidelines for limiting exposure to Electromagnetic Fields (100 kHz to 300 GHz). Health Phys. 2020;118(5):483–524.

35. Muceli S, Poppendieck W, Hoffmann K-P, Dosen S, Benito-León J, Barroso FO, et al. A thin-film multichannel electrode for muscle recording and stimulation in neuroprosthetics applications. J Neural Eng. 2019 Apr 1;16(2):026035.

36. Rose TL, Robblee LS. Electrical stimulation with Pt electrodes. VIII. Electrochemically safe charge injection limits with 0.2 ms pulses (neuronal application). IEEE Trans Biomed Eng. 1990;37(11):1118–20.

37. Poppendieck W, Sossalla A, Krob M-O, Welsch C, Nguyen TAK, Gong W, et al. Development, manufacturing and application of double-sided flexible implantable microelectrodes. Biomed Microdevices [Internet]. 2014;16(6):837–50. Available from: https://doi.org/10.1007/s10544-014-9887-8

38. Muceli S, Poppendieck W, Negro F, Yoshida K, Hoffmann KP, Butler JE, et al. Accurate and representative decoding of the neural drive to muscles in humans with multi-channel intramuscular thin-film electrodes. J Physiol [Internet]. 2015 Sep 1;593(17):3789–804. Available from: http://doi.wiley.com/10.1113/JP270902

39. Poppendieck W, Dörge T, Hoffmann KP. Optimization of microporous platinum coatings for neural microelectrodes. In: 13th Annual International Conference of the IFES Society. 2008. p. 319–21.

40. Meyer J-U, Stieglitz T, Scholz O, Haberer W, Beutel H. High density interconnects and flexible hybrid assemblies for active biomedical implants. IEEE Trans Adv Packag [Internet]. 2001;24(3):366–74. Available from: http://ieeexplore.ieee.org/document/938305/

41. Cogan SF. Neural Stimulation and Recording Electrodes. Annu Rev Biomed Eng. 2008;10(1):275–309.

42. Gabriel C, Gabriel S. Compilation of the Dielectric Properties of Body Tissues at RF and Microwave Frequencies. [Internet]. 1996. Available from: http://niremf.ifac.cnr.it/docs/DIELECTRIC/Report.html

43. Silva A, Campos S, Monteiro J, Venâncio C, Costa B, Guedes de Pinho P, et al. Performance of Anesthetic Depth Indexes in Rabbits under Propofol Anesthesia: Prediction Probabilities and Concentration-effect Relations. Anesthesiology [Internet]. 2011 Aug 1;115(2):303–14. Available from: https://doi.org/10.1097/ALN.0b013e318222ac02

44. Peeters ME, Gil D, Teske E, Eyzenbach V, Brom WE Vd, Lumeij JT, et al. Four methods for general anaesthesia in the rabbit: a comparative study. Lab Anim. 1988;22(4):355–60.

45. Spaich EG, Andersen OK, Arendt-Nielsen L. Tibialis Anterior and Soleus Withdrawal Reflexes Elicited by Electrical Stimulation of the Sole of the Foot during Gait. Neuromodulation Technol Neural Interface [Internet]. 2004 Apr 1;7(2):126–32. Available from: https://doi.org/10.1111/j.1094-7159.2004.04016.x

46. Hasgall P, Di Gennaro F, Baumgartner C, Neufeld E, Lloyd B, Gosselin M, et al. IT’IS Database for thermal and electromagnetic parameters of biological tissues [Internet]. Zurich; 2018. Available from: https://itis.swiss/virtual-population/tissue-properties/

47. Morel P, Ferrea E, Taghizadeh-Sarshouri B, Audí JMC, Ruff R, Hoffmann K-P, et al. Long-term decoding of movement force and direction with a wireless myoelectric implant. J Neural Eng. 2016 Feb 1;13(1):016002.

48. Kampianakis E, Sharma A, Arenas J, Reynold MS. A Dual-Band Wireless Power Transfer and Backscatter Communication Approach for Real-Time Neural/EMG Data Acquisition. IEEE J Radio Freq Identif. 2017;1(1):100–7.

49. McDonnall D, Hiatt S, Smith C, Guillory KS. Implantable multichannel wireless electromyography for prosthesis control. In: 2012 Annual International Conference of the IEEE Engineering in Medicine and Biology Society [Internet]. IEEE; 2012. p. 1350–3. Available from: http://ieeexplore.ieee.org/document/6346188/

50. Weir RF, Troyk PR, DeMichele GA, Kerns DA, Schorsch JF, Maas H. Implantable Myoelectric Sensors (IMESs) for Intramuscular Electromyogram Recording. IEEE Trans Biomed Eng [Internet]. 2009 Jan;56(1):159–71. Available from: http://ieeexplore.ieee.org/document/4633666/

51. Pasquina PF, Evangelista M, Carvalho AJ, Lockhart J, Griffin S, Nanos G, et al. First-in-man demonstration of a fully implanted myoelectric sensors system to control an advanced electromechanical prosthetic hand. J Neurosci Methods [Internet]. 2015 [cited 2017 Jun 7];244:85–93. Available from: http://www.sciencedirect.com/science/article/pii/S0165027014002672

52. Jung MK, Muceli S, Rodrigues C, Megía-García Á, Pascual-Valdunciel A, del-Ama AJ, et al. Intramuscular EMG-Driven Musculoskeletal Modelling: Towards Implanted Muscle Interfacing in Spinal Cord Injury Patients. IEEE Trans Biomed Eng. 2022;69(1):63–74.

53. Salminger S, Sturma A, Hofer C, Evangelista M, Perrin M, Bergmeister KD, et al. Long-term implant of intramuscular sensors and nerve transfers for wireless control of robotic arms in above-elbow amputees. Sci Robot. 2019;4(32):eaaw6306.

54. Ghanbari MM, Piech DK, Shen K, Alamouti SF, Yalcin C, Johnson BC, et al. A Sub-mm3 Ultrasonic Free-Floating Implant for Multi-Mote Neural Recording. IEEE J Solid-State Circuits. 2019;54(11):3017–30.

55. Singer A, Robinson JT. Wireless Power Delivery Techniques for Miniature Implantable Bioelectronics. Adv Healthc Mater. 2021;2100664.

56. Chasset F, Soria A, Moguelet P, Mathian A, Auger Y, Francès C, et al. Contact dermatitis due to ultrasound gel: A case report and published work review. J Dermatol. 2016;43(3):318–20.

57. Becerra-Fajardo L, Garcia-Arnau R, Ivorra A. Injectable Stimulators Based on Rectification of High Frequency Current Bursts: Power Efficiency of 2 mm Thick Prototypes. In: Ibáñez J, González-Vargas J, Azorín JM, Akay M, Pons JL, editors. Converging Clinical and Engineering Research on Neurorehabilitation II. Cham: Springer International Publishing; 2017. p. 667–71.

