## Supplementary file 1 for "Floating EMG Sensors and Stimulators Wirelessly Powered and Operated by Volume Conduction for Networked Neuroprosthetics"

The following sections provide additional details on the methods and results regarding the development and evaluation of the presented system for EMG sensing and electrical stimulation.

#### I. Supplementary Methods

##### 1. External system - communications protocol

Supplementary Table 1 reports the downlink commands included in the communication protocol stack, as well as the estimated time required for them.

Supplementary Table 1
Downlink commands to one or multiple wireless devices

| Function | Description | Transmission time of the frame* | Active time** | Observations |
|---|---|---|---|---|
| Ping | Request ping to a specific wireless device. | 200 µs | 100 µs of 200 µs | Wireless device replies with an ACK. |
| Reset | Request reset to one or a group of wireless devices. | 200 µs | 100 µs of 200 µs | To one device or group. If only one wireless device is requested, it replies with ACK. |
| Set configuration | Configure stimulation. | 280 µs | 140 µs of 280 µs | To one device or group. |
| | Configure sensing. | 280 µs | 140 µs of 280 µs | To one device or group. |
| | Configure group and delivers burst for ACK uplink. | 280 µs | 140 µs of 280 µs | Wireless device replies with an ACK. |
| Get configuration | Requests current configuration of stimulation, and delivers burst to uplink information. | 200 µs | 100 µs of 200 µs | Wireless device replies with current stimulation configuration. |
| | Requests current configuration of sensing, and delivers burst for uplink information. | 200 µs | 100 µs of 200 µs | Wireless device replies with current sensing configuration. |
| | Asks if pertains to a specific group, and delivers burst for ACK uplink. | 280 µs | 140 µs of 280 µs | Wireless device replies with ACK if pertains to that group. |
| Stimulate | Instruction and burst for activating stimulation in wireless device. | 200 µs | 100 µs of 200 µs | To one device or group. External system defines pulse width and frequency of stimulation. Wireless device or group only starts stimulating if the configuration has been set. |
| Start sensing | Requests start sensing. | 200 µs | 100 µs of 200 µs | To one device or group. Wireless device or group only starts sensing if the |

| | | | | |
|---|---|---|---|---|
| | | | | configuration has been set. If only one wireless device is requested, it replies with ACK. |
| Stop sensing | Stop sensing. | 200 µs | 100 µs of 200 µs | To one device or group. If only one wireless device is requested, it replies with ACK. |
| Fast get sample | Requests single acquisition and previous sample, and delivers burst to uplink information. | 120 µs | 60 µs of 120 µs | Does not include identifying code of wireless device as there will be only one device configured as raw in real time. |
| Get sample | Requests sample and delivers burst for corresponding uplink information. | 200 µs | 100 µs of 200 µs | Wireless device replies with a sample. |
| Retry sample | Requests last sample and delivers burst for corresponding uplink information. | 200 µs | 100 µs of 200 µs | In case the sample obtained with "get sample" instruction is not uplinked correctly. |

* To simplify calculations, these values have been rounded up assuming a byte transmission time of 40 µs instead of the actual duration of 39.06 µs (1 start bit + 1 byte + 1 stop bit). The transmission times reported include the synchronization byte time.

** In downlink, active time is 50% of transmission time because of Manchester coding.

The stimulation and sensing configuration payloads of the communication protocol stack are reported in Supplementary Table 2. They can be sent to one wireless device or to a group of devices.

Supplementary Table 2

Stimulation and sensing configuration payloads (downlink)

| Type of configuration | Parameter | Combinations |
|---|---|---|
| Stimulation configuration payload | Type of waveform | Monophasic<br>Biphasic |
| | First pulse | Anodic-first<br>Cathodic-first |
| Sensing configuration payload | Sensing mode | Raw<br>Parametric 1<br>Parametric 2 |
| | Sampling frequency | 250 sps<br>500 sps<br>750 sps<br>1000 sps |
| | Sampling window | Real time<br>Sampling windows (15 options) |

Supplementary Table 3 reports the uplink replies used by the wireless devices to send information to the external system. The replies correspond to requests sent by the external system with a previous downlink command. To do so, the external system waits for 2.3 ms after the downlink command so that the wireless devices can demodulate and decode the information and do further processing to answer the request. Then, the external system delivers a

HF current burst (1 ms) for power maintenance, followed by a 50 µs pause in which no HF current is applied (serves as synchronization flag), a transmission HF burst (timings reported in Supplementary Table 3), and 100 µs power maintenance burst. For example, if an ACK is requested by the external system, the external system delivers a 1 ms burst, followed by a pause of 50 µs, a transmission burst of 80 µs (one synchronization byte and one information byte), and a 100 µs burst, for a total active time of 1.18 ms.

Supplementary Table 3
Uplink replies from one wireless device

| Function | Description | Transmission time of the frame | Total active time |
|---|---|---|---|
| Send ACK | Sends one acknowledgement to the external system. | 80 µs | 1.18 ms of 1.23 ms |
| Send sample | Sends one sample to the external system. | 200 µs | 1.3 ms of 1.35 ms |
| Send configuration | Sends the stimulation configuration to the external system. | 160 µs | 1.26 ms of 1.31 ms |
| | Sends the sensing configuration to the external system. | 160 µs | 1.26 ms of 1.31 ms |

The transmission times reported include the synchronization byte time.

#### 2. Intramuscular electrodes - Design

This section details how the geometry of the electrode contacts of the intramuscular electrodes was determined. In particular, it details how such geometry was designed to 1) ensure that enough power is obtained to supply the floating circuit during the most power consuming mode (i.e., continuous EMG recording), and 2) generate stimulation pulses with amplitudes above 2 mA and below 4 mA.

##### 2.1. Model of tissues surrounding the intramuscular electrode, and electric field

The tissues surrounding the intramuscular electrode and the presence of the electric field applied by the external system can be modeled with a Thévenin equivalent circuit [1]. To obtain the open circuit voltage ($v_{OC}$) and the equivalent impedance ($\boldsymbol{Z}_{Th}$) of that model, it was first performed a finite element method (FEM) study (in COMSOL Multiphysics 4.4) that provided two geometrical scaling factors: $K_{field}$, which translates the electric field magnitude (V/m) into the open circuit voltage across the active sites of the two electrodes; and $K_{Rth}$, which can be simply understood as the scaling factor that transforms the tissue resistivity value (Ω·m) into the equivalent resistance of the Thévenin model. The amplitude ($A$) of the Thévenin voltage source ($v_{OC}$) was scaled as:

$$A = K_{field} \left|\vec{E}\right| \tag{S1}$$

where $\left|\vec{E}\right|$ is the magnitude of the applied electric field. Here, for obtaining $\boldsymbol{Z}_{Th}$, the impedance of tissues was not merely modeled as a resistance but as the parallel combination of the equivalent resistance of the extracellular medium ($R_e$), and the series combination of the equivalent capacitance of the cell membranes ($C_m$), and the equivalent resistance of the intracellular medium ($R_i$). That is, tissues were modeled by a lumped element model with a single Debye relaxation. $R_e$, $R_i$ and $C_m$ were scaled from $K_{Rth}$ as follows:

$$R_{e\_xy} = K_{Rth} \cdot \rho_e [\Omega] \tag{S2}$$

$$R_{i\_xy} = K_{Rth} \cdot \rho_i [\Omega] \tag{S3}$$

$$C_{m\_xy} = K_{field} \cdot c_m [\mathrm{F}] \tag{S4}$$

where $\rho_e$ and $\rho_i$ are the equivalent resistivities of the extracellular and the intracellular media (Ω·m), respectively, and $c_m$ is the volume capacitance (F/m). These three parameters ($\rho_e, \rho_i$ and $c_m$) can be derived from experimental data reported in [2][3]. In the case of muscle tissue their values are: $\rho_e$ = 3.68 Ω·m, $\rho_i$ = 2.84 Ω·m and $c_m$= 0.11 µF/m. The open circuit voltage ($v_{OC}$) and the equivalent impedance ($\boldsymbol{Z}_{Th}$) were incorporated in circuit simulations as described later.

#### 2.2. FEM simulations

The geometry of the FEM simulation consisted in a segment of polyimide filament containing two double-sided active sites (i.e., actual electrodes). The segment had a width of 420 µm, a thickness of 100 µm, and a length of 50 mm (Supplementary Figure 1a). The conductivity of the active sites was set to 1 × $10^6$ S/m and the conductivity of the polyimide substrate was set to 1 × $10^{-5}$ S/m. The segment was simulated within a 0.1 m × 0.1 m × 0.1 m cube with a conductivity of 1 S/m. The segment was centered and aligned with one axis of the cube. Misalignments between the segment and the applied electric field were also simulated ($\alpha$: angle between the electric field and the electrodes' axis).

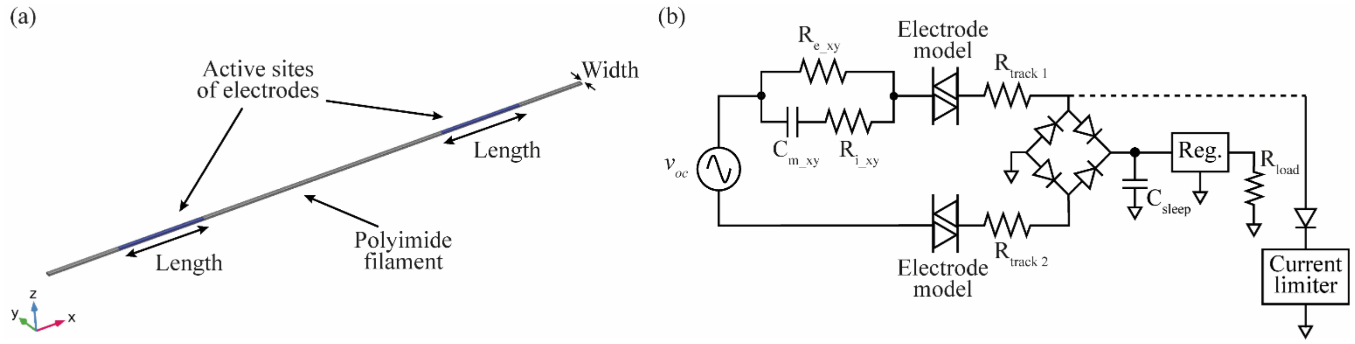


Supplementary Figure 1. Electrode design. (a) Geometry of the intramuscular electrodes for FEM simulation. (b) SPICE circuit that includes the Thévenin equivalent circuit, the electrode model, and the resistance of the tracks that connect the active sites of the electrodes to the circuit's electronics. Two independent simulations were performed: a circuit to assay continuous power supply to a load $R_{load}$, and a circuit to assay the stimulation pulses with a single current limiter (power supply circuit disconnected).

The $K_{field}$ value was obtained by measuring the voltage across the two active sites of the segment when it was simulated the delivery of a 1 V/m electric field. Such field was produced by applying a voltage of 0.1 V across the sides of the cube perpendicular to the segment. The $K_{Rth}$ value was obtained by measuring the voltage across the two active sites of the segment when it was simulated the flow of a current of 1 A through both sites.

Supplementary Table 4 reports the geometrical scaling factors $K_{field}$ and $K_{Rth}$ obtained with the FEM simulation for different geometrical characteristics of the active sites of the electrodes. The distance between the active sites was limited by the maximum length of the polyimide filament, which was limited by the 68 mm diameter of the wafer on which the intramuscular electrodes were fabricated, and the need to ensure that the most proximal active site could be implanted deeply enough, beneath the subcutaneous fat layer.

The width and thickness of the polyimide filament were defined according to the characteristics of previous intramuscular electrodes fabricated using the same technology [4].

Supplementary Table 4
Results from FEM simulations

| Geometrical characteristics | | | Misalignment | Geometrical scaling factors obtained | |
|---|---|---|---|---|---|
| **Width** | **Length** | **Distance** | $\boldsymbol{\alpha}$ | $\boldsymbol{K_{field}}$ | $\boldsymbol{K_{Rth}}$ |
| 200 µm | 4 mm | 30 mm | 0° | 0.0300 m | 294 $m^{-1}$ |
| 200 µm | 4 mm | 30 mm | 30° | 0.0260 m | 294 $m^{-1}$ |
| 200 µm | 7.5 mm | 30 mm | 0° | 0.0300 m | 181 $m^{-1}$ |
| 200 µm | 7.5 mm | 30 mm | 30° | 0.0260 m | 181 $m^{-1}$ |
| 265 µm | 4 mm | 30 mm | 0° | 0.0300 m | 274 $m^{-1}$ |
| 265 µm | 4 mm | 30 mm | 30° | 0.0260 m | 273 $m^{-1}$ |
| 265 µm | 7.5 mm | 30 mm | 0° | 0.0300 m | 171 $m^{-1}$ |
| 265 µm | 7.5 mm | 30 mm | 30° | 0.0263 m | 171 $m^{-1}$ |
| 380 µm | 2 mm | 20 mm | 0° | 0.0200 m | 394 $m^{-1}$ |
| 380 µm | 2 mm | 20 mm | 30° | 0.0167 m | 394 $m^{-1}$ |
| 380 µm | 4 mm | 30 mm | 0° | 0.0303 m | 244 $m^{-1}$ |
| 380 µm | 4 mm | 30 mm | 30° | 0.0259 m | 247 $m^{-1}$ |
| 380 µm | 5 mm | 30 mm | 0° | 0.0292 m | 210 $m^{-1}$ |
| 380 µm | 5 mm | 30 mm | 30° | 0.0259 m | 210 $m^{-1}$ |
| 380 µm | 5 mm | 40 mm | 0° | 0.0395 m | 212 $m^{-1}$ |
| 380 µm | 5 mm | 40 mm | 30° | 0.0356 m | 212 $m^{-1}$ |

FEM simulations indicate that the value of the geometrical scaling factor $K_{field}$, which is proportional to the Thévenin voltage source of the Thévenin model (Supplementary Figure 1b), is approximately equal to

$$K_{field} = \cos(\alpha) \times L \quad \text{(S5)}$$

where $\alpha$ is the misalignment angle, and $L$ is the length of the electrode contact. It is quite likely that the correct value is that obtained analytically, and that the small discrepancies between the expression and the simulation results (Supplementary Table 4) are due to numerical errors during the simulations.

#### 2.3. SPICE simulations

The two geometrical scaling factors ($K_{field}$ and $K_{Rth}$) were used in SPICE simulations (LTspice XVII by Analog Devices, Inc.). The simulations include a resistance ($R_{track}$) for modeling the resistance of the tracks that electrically connect the active sites to the pads where the floating circuit is connected; and the so-called electrode/electrolyte impedance with a model extracted from literature [5]. Two circuits were simulated: a circuit to assay continuous power supply, and a circuit to assay the stimulation pulses (Supplementary Figure 1b). For the former, a capacitor ($C_{sleep}$) is used to smooth the input of a 2.5 V ideal regulator that supplies a resistive load ($R_{load}$) with a current of 300 µA. For the latter, a realistic model for the electrical stimulation subcircuit implemented in the floating devices was simulated. The architecture of the subcircuit is explained below. The maximum values for stimulation amplitude, pulse duration and repetition frequency were set at 4 mA, 400 µs and 100 Hz respectively, while the power supply circuit was disconnected. In both simulations, the applied electric field was calculated by defining a SAR of 2 W/kg, using bursts with a frequency (*F*) of 50 Hz, and a duration (*B*) of 1.6 ms.

The obtained geometrical scaling factors reported in Supplementary Table 4 were assayed in the two SPICE simulations proposed. Here are reported the results for the final conformation, in which the active site has a width of 265 µm and a length of 7.5 mm, and the active sites have a separation distance of 30 mm. With this geometry, the calculated resistance of the distal ($R_{track1}$) and proximal ($R_{track2}$) tracks is 49.1 Ω and 25.6 Ω respectively.

#### 3. Electrical stimulation subcircuit of the miniature electronic circuit

Electrical stimulation is performed using two independent current limiters, each one connected to a Schottky diode (RB521ZS-30 by ROHM Co., Ltd.)  that is connected to the dc-blocking capacitor shown in Fig. 5 (e). When the control unit of the floating device identifies the stimulation bursts, it activates a specific current limiter depending on the polarity set during configuration. Supplementary Figure 2 shows an example in which biphasic cathodic-first stimulation is done. When current limiter 1 (yellow) is activated by the control unit, it forces the flow of stimulating (half-wave) rectified current through it, generating a negative low-frequency current seen from the tissues. When the second current limiter is activated, it forces the flow of current to go in the opposite direction, generating a positive low-frequency current seen from the tissues.

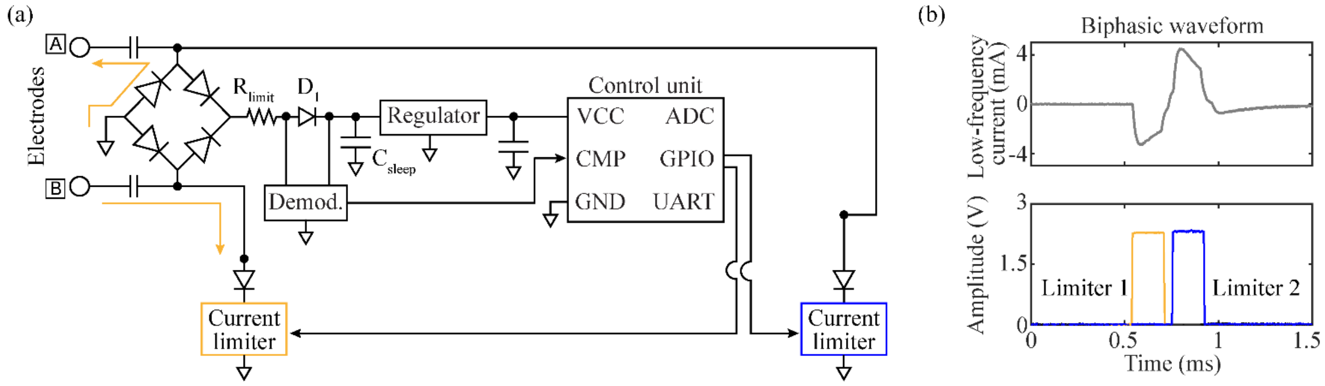


Supplementary Figure 2. Example of sequence for stimulation, and how the current limiter forces the flow of current in one direction or the other. (a) Basic circuit architecture showing only the subcircuits required in the electrical stimulation mode. (b) Corresponding results of filtered low-frequency current applied, and activation of current limiters.

The architecture of the current limiter is shown in Supplementary Figure 3. A BJT NPN transistor (BC847BFZ by Diodes Incorporated) acts as an output transistor ($Q_1$), and a second identical transistor ($Q_2$) acts as a protection transistor. When the voltage across the sensing resistor ($R_{sense}$) is lower than the base-emitter voltage of $Q_2$ ($V_{BE}$), only $Q_1$ is active, and the current on the load ($I_{load}$) will increase. As soon as the voltage across $R_{sense}$ is higher than

$V_{BE}$, $Q_2$ turns on and draws the base current of $Q_1$, reducing its collector current, therefore limiting $I_{load}$. Using this topology, the current flowing through $R_{sense}$ (i.e., $I_{load}$) depends on the value of this resistor. Each current limiter is connected/disconnected from the load (i.e., the tissue) using a switch based on a N-channel MOSFET (DMN2990UFZ by Diodes Incorporated) controlled by the control unit of the floating device using one GPIO.

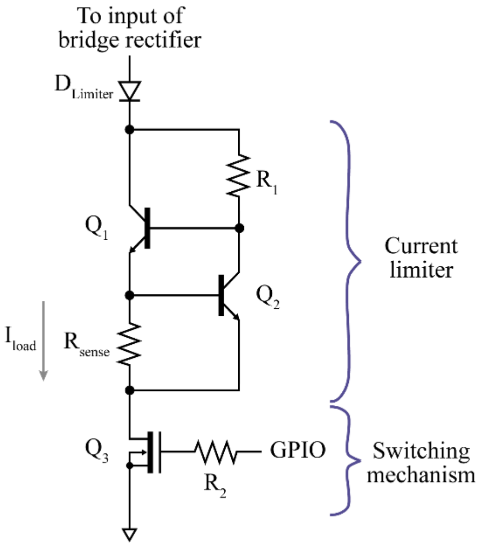


Supplementary Figure 3. Architecture of the current limiter's subcircuit.

### II. Supplementary Results

#### 1. SPICE simulations

To test the ability of the basic circuit (i.e., bridge rectifier, $C_{sleep}$, low-dropout linear regulator, a resistor as a load, and a current limiter) to obtain a stable power supply using the geometrical scaling factors reported above, the simulated voltage source of the external system was configured to deliver an initial Power up burst of 30 ms, followed by short bursts for power maintenance (*F*: 50 Hz; *B*: 1.6 ms). Supplementary Figure 4 shows that the circuit is able to obtain a steady output at the regulator after 350 ms from the start of the Power up. When the power maintenance bursts are delivered by the external system (Supplementary Figure 4, top), $C_{sleep}$ is charged and the voltage across it increases accordingly (label "Regulator in").

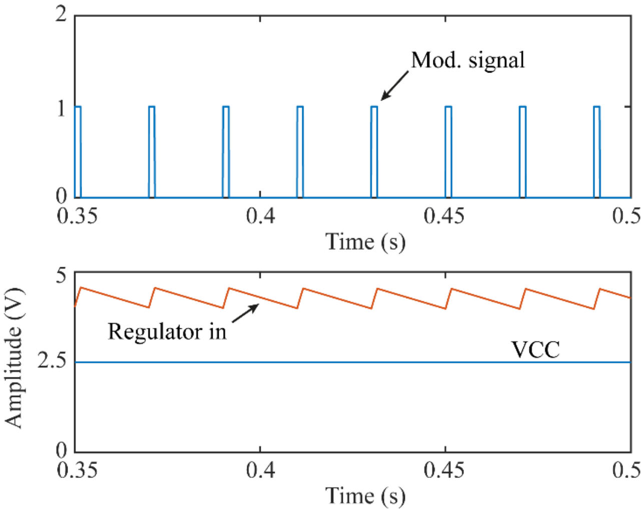


Supplementary Figure 4. SPICE simulation result of continuous power supply using the geometrical scaling factors obtained for a width of 265 µm, a length of 7.5 mm, and a separation distance of 30 mm (no misalignment). Top: Modulating signal corresponding to the times when the short bursts are delivered by the voltage source. Bottom: Electric potential difference seen at the regulator's input and output (VCC).

Supplementary Figure 5 shows the low frequency current delivered by the simulated circuit when the current limiter is activated. In this particular example, the current limiter is activated to deliver stimulation pulses with a pulse width of 400 µs, at a frequency of 100 Hz. This current is measured using a virtual LPF (cutoff frequency: 10 kHz) of the current flowing through the track resistor $R_{track2}$. According to simulations, the intramuscular electrodes can deliver current with amplitudes above 4 mA to perform electrical stimulation.

The proposed width (0.260 mm) and length (7.5 mm) for each active site creates a total area of 3.9 $mm^2$ for each electrode (two-sided electrode). Assuming stimulation pulses of maximum 400 µs, and a very conservative maximum charge injection capacity of 50 $\mu C/cm^2$, the circuit should deliver a maximum current of 4.9 mA to avoid irreversible reactions in the electrodes. This limit is above the maximum current required in the application proposed here (4 mA).

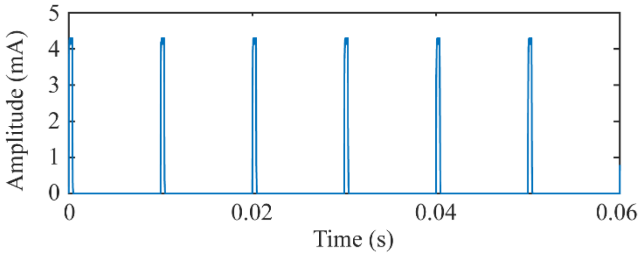


Supplementary Figure 5. SPICE simulation result of stimulation pulses delivered by the intramuscular electrodes using the geometrical scaling factors obtained for a width of 265 µm, a length of 7.5 mm, and a separation distance of 30 mm (no misalignment).

#### 2. Compliance with electrical safety standards

The compliance with electrical safety standards study is based in the *in vitro* scenario. In this study the external system was set to apply HF current bursts with a frequency of 3 MHz and an amplitude of 39 V, equivalent to a peak electric field of 325 V/m. Supplementary Table 5 reports the calculated duty cycle (3), $E_{rms}$ (2) and SAR calculated at a point (1) averaged over 6 minutes ($\sigma$: 0.57 S/m [6] and $\rho$: 1090 kg/m$^3$ [7] for muscle tissue at 3 MHz). using three different sequences. Sequence A indicates a sequence in which a Power up is applied, followed by power maintenance bursts. Sequence B proposes a sequence in which there is a Power up, followed by the instructions required to configure stimulation (biphasic, cathodic-first), and then 10,000 biphasic pulses (200 Hz, 400 µs pulse width, 30 µs interphase dwell) corresponding to 50 seconds of stimulation, and the reminder time used for power maintenance. Sequence C exemplifies a sequence in which a Power up is applied, followed by the configuration of EMG acquisition (raw, 500 sps), configuration of stimulation (biphasic, cathodic-first) followed by 120 cycles with a duration of 3 s, and that include: 1 s of EMG sensing (i.e., Start sensing is sent, followed by a Stop sensing 1 s after), 1 s of samples uplink and external processing (i.e., 500 Get samples followed by power maintenance bursts while the external system processes the results), and 1 s of stimulation (i.e., 100 biphasic pulses at 100 Hz, 200 µs pulse width, 30 µs interphase dwell).

Supplementary Table 5
Duty cycle, $E_{rms}$ and SAR calculated for different bidirectional sequences with an averaging time of 6 minutes

| **Sequence** | **Duty cycle** | **$E_{rms}$ (V/m)** | **SAR (W/kg)** |
|---|---|---|---|
| A. Power and maintenance | 0.08 | 65.03 | 2.21 |
| B. Power, config. Stimulation, stimulate, and maintenance | 0.12 | 80.63 | 3.40 |
| C. Power, config., 120 cycles of sensing, samples uplink, and stimulation; and maintenance | 0.29 | 124.44 | 8.10 |

#### 3. EMG sensors reported in the literature

Supplementary Table 6 compares different implantable EMG sensors reported in the literature.

Supplementary Table 6

Comparison of implantable EMG sensors reported in literature

| Name | Powering method | Electrode type | Gain | Bandwidth | ADC resolution | Form factor |
|---|---|---|---|---|---|---|
| Farnsworth et al. [8] | Inductive coupling | Epimysial | 38 dB | 1000 Hz | 11 bits | Central unit with leads; 6 mm Ø |
| IMES [9] [10] | Inductive coupling | Intramuscular | 24.1 – 78 dB* | 1000 Hz | 8 bits | Cylindrical; 2.5 mm Ø, 16 mm long |
| IST-12 [11] | Inductive coupling | Epimysial | 46 – 78 dB* | 900 Hz | 12 bits | Central unit with leads; 40 × 38 × 7 mm (dimensions of electronics only) |
| MyoPlant [12][13] | Inductive coupling | Epimysial | 33.6 –61 dB* | 1500 Hz | 10 bits | Central unit with leads, 38 × 25 × 8 mm |
| Ripple [14] | Inductive coupling | Epimysial and intramuscular | 46 dB | NA | 12 bits | Central unit with leads; 70 × 35 mm |
| IEAD [15,16] | Inductive coupling | Intramuscular | 55, 61.6 or 77.1 dB* | 5.0, 5.5 or 7.3 kHz | 10 bits | Disc; 18 mm Ø, and central unit with leads; 10 × 20 mm |
| Reynolds et al. [17,18] | Inductive coupling | NA | 34 dB | 700 Hz | 11 bits | Central unit with leads; 25 mm Ø, 2.8 mm thick |
| This work | Volume conduction | Intramuscular | 54 dB | 1000 Hz | 10 bits | |

* Programmable
NA: information not available
Ø: diameter

#### References


1. Tudela-Pi M, Becerra-Fajardo L, García-Moreno A, Minguillon J, Ivorra A. Power Transfer by Volume Conduction: In Vitro Validated Analytical Models Predict DC Powers above 1 mW in Injectable Implants. IEEE Access. 2020;1.
2. Andreuccetti D, Fossi R, Petrucci C. An Internet resource for the calculation of the dielectric properties of body tissues in the frequency range 10 Hz - 100 GHz. Website at http://niremf.ifac.cnr.it/tissprop/. IFAC-CNR, Florence (Italy), 1997. Based on data published by C.Gabriel et al. in 1996. 1997.
3. Gabriel S, Lau RW, Gabriel C. The dielectric properties of biological tissues: III. Parametric models for the dielectric spectrum of tissues. Phys Med Biol. 1996;41(11):2271–93.
4. Muceli S, Poppendieck W, Hoffmann K-P, Dosen S, Benito-León J, Barroso FO, et al. A thin-film multichannel electrode for muscle recording and stimulation in neuroprosthetics applications. J Neural Eng. 2019 Apr 1;16(2):026035.
5. Jones MH, Scott J. Scaling of Electrode-Electrolyte Interface Model Parameters In Phosphate Buffered Saline. IEEE Trans Biomed Circuits Syst. 2015;9(3):441–8.

6. Gabriel C, Gabriel S. Compilation of the Dielectric Properties of Body Tissues at RF and Microwave Frequencies. [Internet]. 1996. Available from: http://niremf.ifac.cnr.it/docs/DIELECTRIC/Report.html
7. Hasgall P, Di Gennaro F, Baumgartner C, Neufeld E, Lloyd B, Gosselin M, et al. IT'IS Database for thermal and electromagnetic parameters of biological tissues [Internet]. Zurich; 2018. Available from: https://itis.swiss/virtual-population/tissue-properties/
8. Farnsworth BD, Triolo RJ, Young DJ. Wireless implantable EMG sensing microsystem. In: 2008 IEEE Sensors [Internet]. IEEE; 2008 [cited 2017 Sep 8]. p. 1245–8. Available from: http://ieeexplore.ieee.org/document/4716669/
9. Weir RF, Troyk PR, DeMichele GA, Kerns DA, Schorsch JF, Maas H. Implantable Myoelectric Sensors (IMESs) for Intramuscular Electromyogram Recording. IEEE Trans Biomed Eng [Internet]. 2009 Jan;56(1):159–71. Available from: http://ieeexplore.ieee.org/document/4633666/
10. Salminger S, Sturma A, Hofer C, Evangelista M, Perrin M, Bergmeister KD, et al. Long-term implant of intramuscular sensors and nerve transfers for wireless control of robotic arms in above-elbow amputees. Sci Robot. 2019;4(32):eaaw6306.
11. Hart RL, Bhadra N, Montague FW, Kilgore KL, Peckham PH. Design and Testing of an Advanced Implantable Neuroprosthesis With Myoelectric Control. IEEE Trans Neural Syst Rehabil Eng [Internet]. 2011 Feb [cited 2018 Apr 18];19(1):45–53. Available from: http://ieeexplore.ieee.org/document/5585827/
12. Morel P, Ferrea E, Taghizadeh-Sarshouri B, Audí JMC, Ruff R, Hoffmann K-P, et al. Long-term decoding of movement force and direction with a wireless myoelectric implant. J Neural Eng. 2016 Feb 1;13(1):016002.
13. Lewis S, Russold M, Dietl H, Ruff R, Audí JMC, Hoffmann K-P, et al. Fully Implantable Multi-Channel Measurement System for Acquisition of Muscle Activity. IEEE Trans Instrum Meas. 2013;62(7):1972–81.
14. McDonnall D, Hiatt S, Smith C, Guillory KS. Implantable multichannel wireless electromyography for prosthesis control. In: 2012 Annual International Conference of the IEEE Engineering in Medicine and Biology Society [Internet]. IEEE; 2012. p. 1350–3. Available from: http://ieeexplore.ieee.org/document/6346188/
15. Ng KA, Rusly A, Gammad GGL, Le N, Liu S-C, Leong K-W, et al. A 3-Mbps, 802.11g-Based EMG Recording System With Fully Implantable 5-Electrode EMGxbrk Acquisition Device. IEEE Trans Biomed

Circuits Syst. 2020;14(4):889–902.

16. Ng KA, Yuan C, Rusly A, Do A-T, Zhao B, Liu S-C, et al. A Wireless Multi-Channel Peripheral Nerve Signal Acquisition System-on-Chip. IEEE J Solid-State Circuits. 2019;54(8):2266–80.

17. Kampianakis E, Sharma A, Arenas J, Reynold MS. A Dual-Band Wireless Power Transfer and Backscatter Communication Approach for Real-Time Neural/EMG Data Acquisition. IEEE J Radio Freq Identif. 2017;1(1):100–7.

18. Thomas SJ, Harrison RR, Leonardo A, Reynolds MS. A Battery-Free Multichannel Digital Neural/EMG Telemetry System for Flying Insects. IEEE Trans Biomed Circuits Syst. 2012;6(5):424–36.
